# Lung macrophages mediate helminth resistance through differential activation of recruited monocyte-derived alveolar macrophages and arginine depletion

**DOI:** 10.1101/2020.10.02.322388

**Authors:** Fei Chen, Darine W. El-Naccache, John J. Ponessa, Alexander Lemenze, Vanessa Espinosa, Wenhui Wu, Katherine Lothstein, Linhua Jin, Olivia Antao, Jason S. Weinstein, Payal Damani-Yokota, Kamal Khanna, Peter Murray, Amariliz Rivera, Mark C. Siracusa, William C. Gause

## Abstract

Macrophages are known to mediate anti-helminth responses, but it remains uncertain which subsets are involved or how macrophages actually kill helminths. Here we show rapid monocyte recruitment to the lung after infection with the nematode parasite, *Nippostrongylus brasiliensis*. In this inflamed tissue microenvironment these monocytes differentiate into an alveolar-like macrophage (AM) phenotype, expressing both Siglec-F and CD11c, surround invading parasitic larvae and preferentially kill parasites in vitro. Monocyte-derived AMs (Mo-AMs) express type 2-associated markers and show distinct remodeling of the chromatin landscape relative to tissue-derived AMs. In particular, they express high amounts of Arg1 (arginase-1), which we demonstrate mediates helminth killing through L-arginine depletion. These studies indicate that recruited monocytes are selectively programmed in the pulmonary environment to express AM markers and an anti-helminth phenotype.

## Introduction

Helminth infections trigger potent type 2 immune responses characterized by elevations in IL-4, IL-5, and IL-13. This response mediates host protection through limiting parasite burden and by mitigating tissue damage associated with the trafficking of these large multicellular parasites through host tissues^1^. Although helminth infections represent a major global health problem, effective vaccine strategies remain elusive, likely in part because the cellular and molecular mechanisms through which the anti-helminth immunity is mediated remain unclear. A critical stage of the life cycle of many helminths, including intestinal nematode parasites, involves transit through the host lung. Previous studies have indicated that the lung plays a significant role in early parasite clearance and as such is a potential target for therapeutic and vaccine-strategies to promote host resistance ^2, 3^.

Both lymphoid and myeloid cells play important roles in promoting type 2 inflammation and anti-helminth immunity. For example, T cells, B cells, myeloid cells and innate lymphoid type 2 cells are activated and promote robust levels of IL-4, IL-5 and IL-13 following a helminth challenge. Importantly, studies also indicate that macrophages and neutrophils contribute to host protection by directly killing parasitic larvae and initiating the healing of parasite affected tissues^3–6^. Neutrophils not only directly target helminths^4^, but also stimulate macrophages to assume an alternatively activated state required for their anti-helminth effector functions^3, 7^. Recent studies indicate that macrophages primed by helminths in the intestine, lung and skin can mediate helminth killing both in vitro and in these tissue microenvironments^2, 3, 8–11^.

Tissue-resident macrophage populations can be subdivided into tissue-derived macrophages (TD-macs) and monocyte-derived macrophages (Mo-macs). TD-macs develop during the early stages of fetal development from the yolk sac and fetal liver, and populate tissues under steady-state conditions, whereas Mo-macs develop from bone marrow-resident precursor cells and can enter tissues in the context of inflammation^12–15^. In the lung microenvironment, tissue-derived alveolar macrophages (TD-AMs) mediate homeostasis through multiple mechanisms, including surfactant recycling, and also provide surveillance for early recognition of pathogens invading the airways^16^. Other scarce tissue-derived interstitial macrophage populations also can have distinct properties^17^. During infection monocytes are rapidly recruited to the lung, where they can differentiate into macrophages. Recent studies suggest that monocytes can also differentiate into an alveolar macrophage-like phenotype and that these Mo-AMs can function differently than TD-AMs^18^. In one study, the Mo-AMs were the predominate lung macrophage subset contributing to fibrosis in a bleomycin-induced lung injury model^19^. Despite these substantial advances in understanding macrophage ontogeny, studies examining the development of Mo-AMs have primarily relied on radiation chimeras or other non-physiological approaches raising questions as to the physiological significance of these observations. Further, whether Mo-AMs populate the lung and perform distinct functions in the context of helminth infections remains to be determined.

Although the anti-helminth qualities of macrophages are well-documented, there is a substantial knowledge gap regarding the mechanisms they employ to promote parasite killing. Previous studies have shown that Arginase (Arg) activity^3, 8, 20^, specifically Arg 1^11^, is required for the anti-helminth properties of macrophages. Arg1 is known to mediate a cascade of effects leading to potentially toxic urea production and also polyamine synthesis which may contribute to immune cell activation and proliferation^20^. Arg1 can also mediate the localized depletion of arginine, which can have immunoregulatory effects on activated immune cells such as T cells^21^. However, whether any of these or other potentially associated mechanisms are important in mediating macrophage killing of helminth parasites and whether they are distinctly initiated in specific subsets of macrophages remains uncertain.

Infection with the intestinal nematode parasite, *N. brasiliensis* (Nb) is a well-established rodent experimental model showing a similar life cycle to human hookworms. Use of this model has substantially advanced our understanding of inflammation and macrophage biology in lung damage and repair caused by transiting helminths. When infected with Nb, infective third-stage larvae (L3) migrate from the skin to the lung before entering the small intestine. In the lung, both innate and adaptive components of the type 2 immune response interact with the migrating parasitic larvae^2, 5, 6^. Previous studies have shown that CD4 T cell depletion at the time of secondary inoculation does not affect accelerated resistance, which is also intact in B cell deficient mice^22, 23^. However, lung macrophages primed by primary Nb inoculation have previously been shown to play an important role in acquired resistance and highly purified helminth-primed lung macrophages can effectively kill L3 in vitro^3^.

Here we investigated the role of lung macrophage subsets of different progenitor origins in mediating acquired resistance to Nb. We further elucidate the role of neutrophils in driving their effector function and reveal how macrophages mediate resistance against helminths via Arg1-dependent mechanisms.

## Results

Nb L3 enter the lung as early as 12 hours after subcutaneous inoculation. To examine how parasite migration through the lung affected numbers of monocytes and macrophages, these populations were analyzed via flow cytometric analysis on days 1, 7, and 14 after inoculation. Marked increases in the total number of macrophages (F4/80^+^, CD64^+^) were observed by day 7 after inoculation (**Fig. 1a**) while monocytes (F4/80^+,^ Ly6C^+^) were markedly increased within 24 hours, consistent with their rapid recruitment, peaked at day 7, and had dropped markedly by day 14 (**Fig. 1b**). Alveolar macrophages (AMs) (CD64^+^,F4/80^+^,SiglecF^+^, CD11c^+^) were slightly elevated as early as day 1 but showed marked increases, more than doubling by day 7 (**Fig. 1c**). Non alveolar macrophages (hereafter, referred to as NAMs) (CD64^+^, F4/80^+^, Siglec-F^-^, CD11c^var^), were also markedly elevated by day 7 after Nb inoculation (**Fig. 1d**). In comparison, eosinophils, also associated with Nb infection, were not elevated until day 7 after inoculation and remained elevated at day 14 (**Fig. 1e**). Collectively, these data demonstrate that Nb infection promotes substantial changes in lung monocyte and macrophage subset dynamics. Further, these data suggest that the early influx of monocytes observed in the lung may help to support the overall increases in lung macrophage populations observed post-infection.

**Fig. 1.**
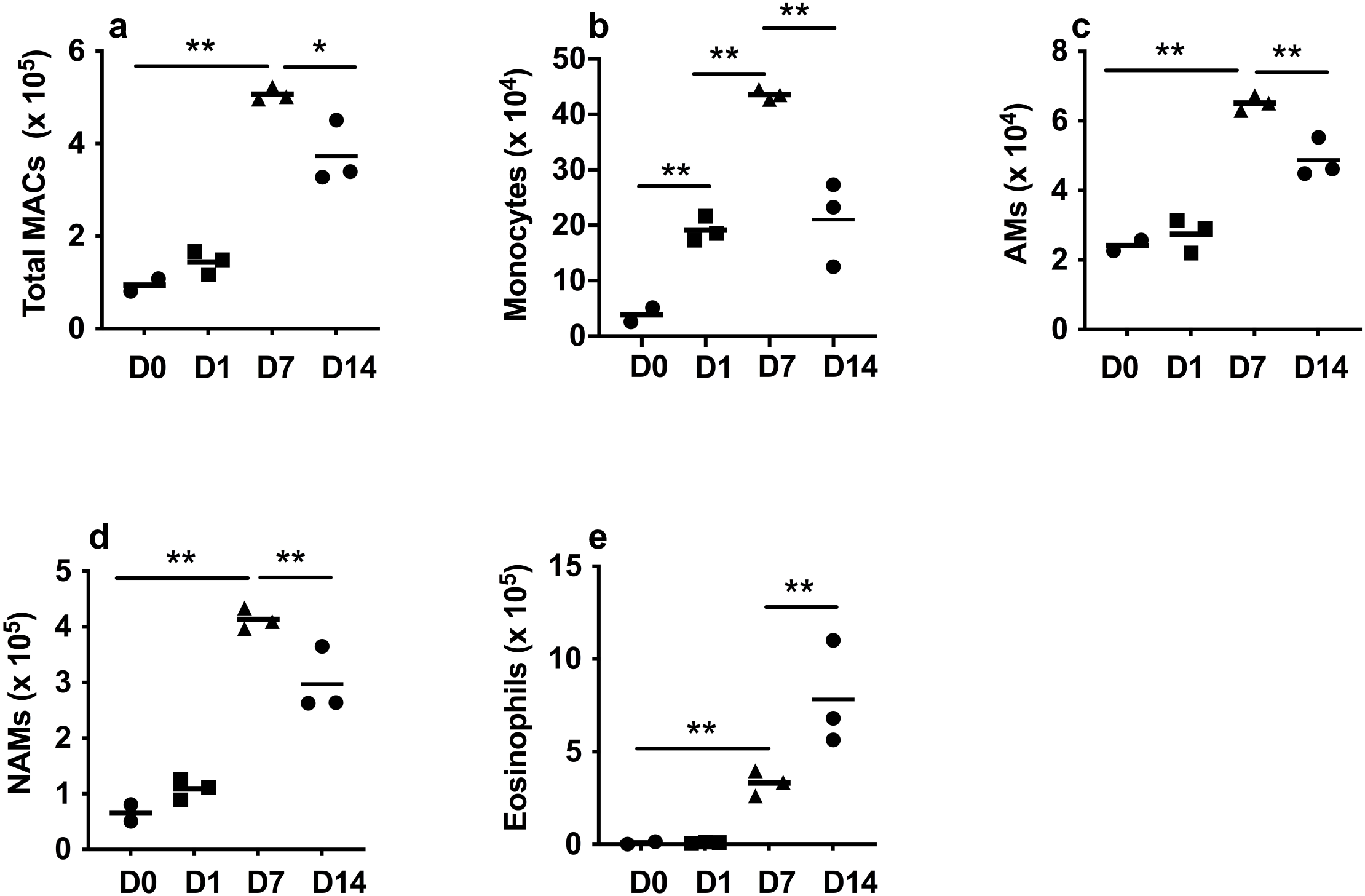
Substantial changes in lung macrophage subsets occurs after *N. brasiliensis* inoculation. Flow cytometric analysis of macrophage subsets obtained from single cell suspensions of whole lungs from BALB/c mice at days 1, 7, and 14 after *N. brasiliensis* inoculation. (a) Total numbers of lung macrophages (CD64^+^, F4/80^+^, CD11c^var^); (b) monocytes (CD11b^+^,Ly6C^+^); (c) alveolar macrophages (AMs), (CD64+,F4/80+,SiglecF+,CD11c+); (d) non-alveolar macrophages (NAMs) (CD64^+^,F4/80^+^,Siglec-F^-^,CD11c^var^); (e) eosinophils (F4/80^+^,Siglec-F^+^, CD11c^-^). Each symbol represents an individual mouse and horizontal lines indicate the mean. All results are representative of two independent experiments. *p<0.05, **p<0.01.

Our previous studies have shown that by day 7 after Nb inoculation sort-purified lung macrophages can effectively kill infectious Nb larvae (L3) following in vitro culture and can mediate accelerated resistance when transferred to otherwise naïve recipients^3^. Importantly, these bulk macrophage populations are comprised of both AMs and NAMs. However, whether these two macrophage subsets equally or preferentially contribute to parasite killing remains unknown. To test this, at day 7 after Nb inoculation, lung cell suspensions were prepared as previously described^3^, and AMs and NAMs were sort-purified. Individual macrophage subsets (1x10^6^/ml) were then cultured with exsheathed L3 (100/well) in 24 well plates, for 5 days, as previously done with bulk macrophages^3^. As shown in **Fig. 2a**, AMs were approximately 5 times more effective at killing L3 than NAMs. Assessment of parasite metabolic activity, by assaying their ATP amounts^3^, further showed significantly decreased activity in cultures seeded with AMs compared to cultures seeded with NAMs (**Fig. 2b**). Previous studies have shown that Arg1 production by macrophages is essential for the effective killing of parasitic larvae both in vitro and in vivo^6, 8^. Therefore, to investigate whether the macrophage killing observed correlated with Arg1 production, AMs and NAMs were assessed for cytoplasmic Arg1 expression at 7 days after Nb infection. As shown in **Fig. 2c**, more than 10% of AMs expressed Arg1 after infection, while less than 3% of NAMs from treated mice expressed Arg1, consistent with NAMs being less effective in killing L3. Furthermore, the total number of Arg1 expressing AMs was significantly greater post-infection while the numbers of Arg1-expressing NAMs remained unchanged (**Fig. 2d**). Collectively, these studies indicate that AMs mediate the majority of the parasite killing capacity within lung macrophage compartments and that elevated cytoplasmic Arg1 protein expression is associated with that enhanced capacity in a subset representing ∼10% of the total AM compartment.

**Fig. 2.**
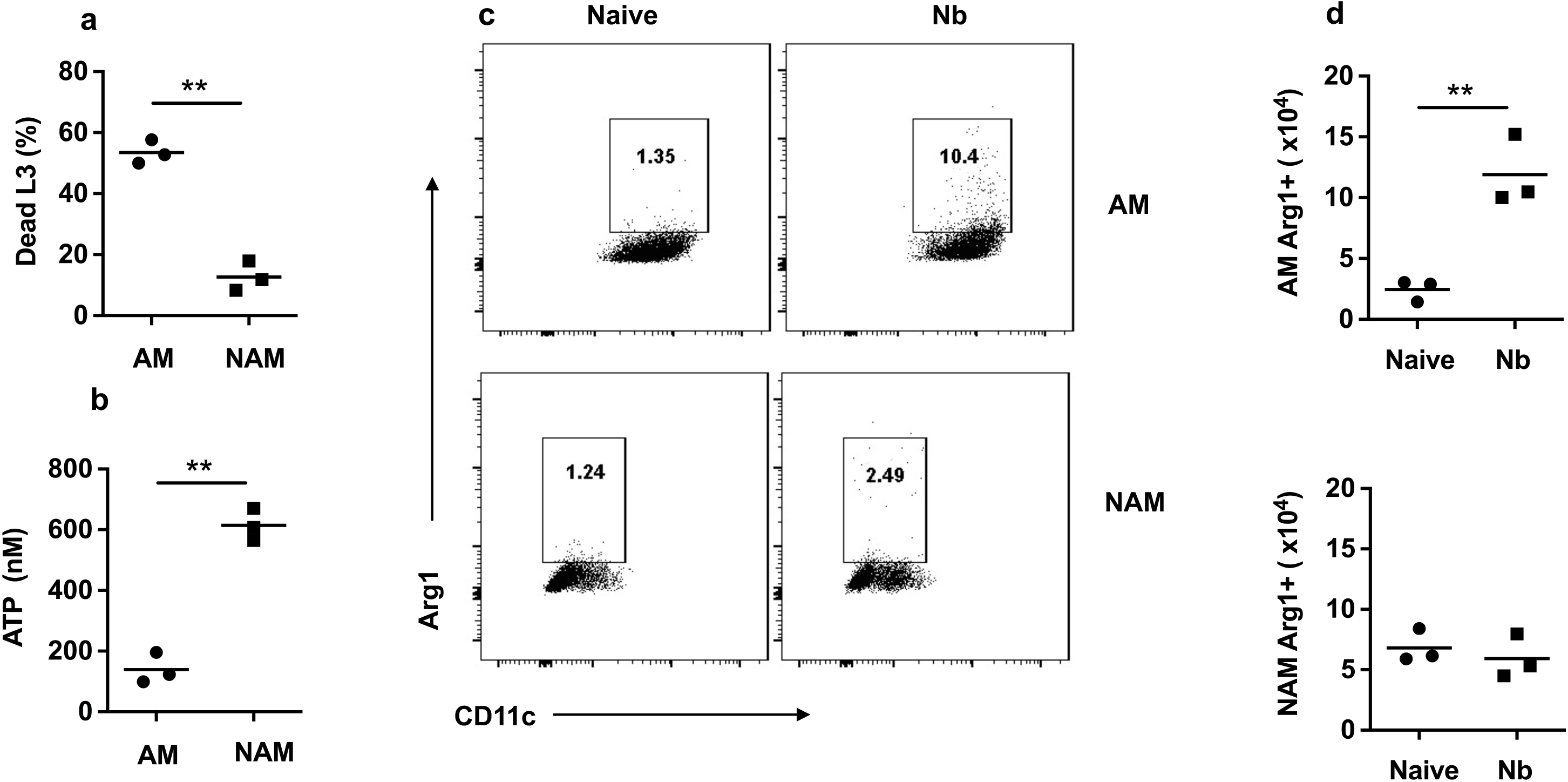
Alveolar macrophages preferentially contribute to parasite killing and express high arginase. (a, b) Lung alveolar macrophages (AMs) and nonalveolar macrophages (NAMs) were stained as in Fig. 1 and sort purified from whole lung cell suspensions at day 7 after *N. brasiliensis* inoculation of BALB/c mice. AMs and NAMs were seeded to 24-well plates (1x 10^6^ cell/well) and co-cultured with 100 exsheathed third stage larvae (L3). At day 5 after culture, the percent mortality (a) and worm ATP concentration (b) were determined. This experiment was repeated three times with similar results. (c, d) Representative flow cytometry plots (c) and lung macrophage numbers (d) for cytoplasmic arginase protein expression in AMs and NAMs at day 7 after *N. brasiliensis* inoculation. Each symbol represents triplicates of pooled samples (a, b), or individual mice (d). All results are representative of two independent experiments. **p<0.01

Our previous studies showed that the transfer of bulk lung macrophages from Nb primed mice into otherwise naïve recipients was sufficient to mediate accelerated resistance similar to the memory response observed after secondary challenge^3^. To examine whether AMs also showed preferential resistance in vivo, AMs and NAMs were sort-purified from the lungs of donor mice at day 7 after Nb inoculation and transferred to naïve recipient mice, which were then inoculated with Nb. Unexpectedly, as shown in **Fig. 3a**, both AMs and NAMs were equally effective at mediating accelerated acquired resistance in recipient mice at 5 days after Nb inoculation. Given that the NAMs expressed low Arg1 relative to the AMs, this raised the possibility that the transferred NAMs were differentiating into an AM-like subset that acquired the ability to preferentially upregulate Arg1 and kill invading parasitic larvae. To test whether NAMs could develop an AM-like phenotype in the lung, we isolated AMs and NAMs from CD45.1 donor mice at day 7 after Nb inoculation and transferred these subsets intratracheally to naïve congenic CD45.2 recipients. Recipient mice were rested 2 days, inoculated with Nb and 5 days later lung cell suspensions were collected and assessed for donor and recipient AM populations. As shown in **Fig. 3b**, transferred CD45.1 donor NAMs gave rise to a substantial number of macrophages with an AM-like phenotype (CD45.1^+^, CD64^+^,F4/80^+^,SiglecF^+^, CD11c^+^) that represented about 10% of the total AM population in the CD45.2 congenic recipients. These data suggest that monocyte-derived macrophages contained within the NAM compartment may possess the capacity to acquire an AM-like phenotype. To further investigate the contribution of monocytes to the AM compartment post-Nb, we obtained CCR2 GFP reporter and CCR2 Diphtheria toxin receptor (DTR) depleter mice. As early as day 1 after Nb inoculation, CCR2^+^ monocytes were increased several fold in the lung and these high levels were sustained as late as day 7 after Nb inoculation (**Fig. 3c**). These data further support our findings presented in **Fig. 1b** and highlight the significant recruitment of monocytes into the lung shortly after Nb inoculation. To further examine the extent to which monocytes contributed to the overall lung macrophage population after infection, CCR2^DTR^ mice were inoculated with Nb and administered DT at -1, 1, and 3 days post-infection to deplete monocytes and monocyte-derived macrophages. As expected, NAMs were significantly reduced on day 7 post-infection after DT administration (**Fig. 3d**), further corroborating a substantial contribution of monocytes to the NAM compartment. However, intriguingly, AMs also showed a marked reduction in their expansion typically observed after Nb inoculation (**Fig. 3e**). These data are consistent with a role for monocyte-derived cells contributing to the population expansion of AMs post-infection (**Fig. 3e**). To confirm the selective depletion of DT treatment on CCR2^+^ monocyte-derived macrophages post-DT treatment, a separate myeloid population, eosinophils, which are CCR2^-^, were also assessed. Eosinophils were also increased in the lung at 5 days after inoculation but were not affected by DT administration consistent with the specificity of the treatment (**Fig. 3f**). In aggregate, these data suggest that lung infiltrating monocytes can acquire a tissue-resident AM phenotype and contribute substantially to the increased AM population observed post-infection. However, the contributions of these monocyte-derived AMs to host protection remained unknown.

**Fig. 3.**
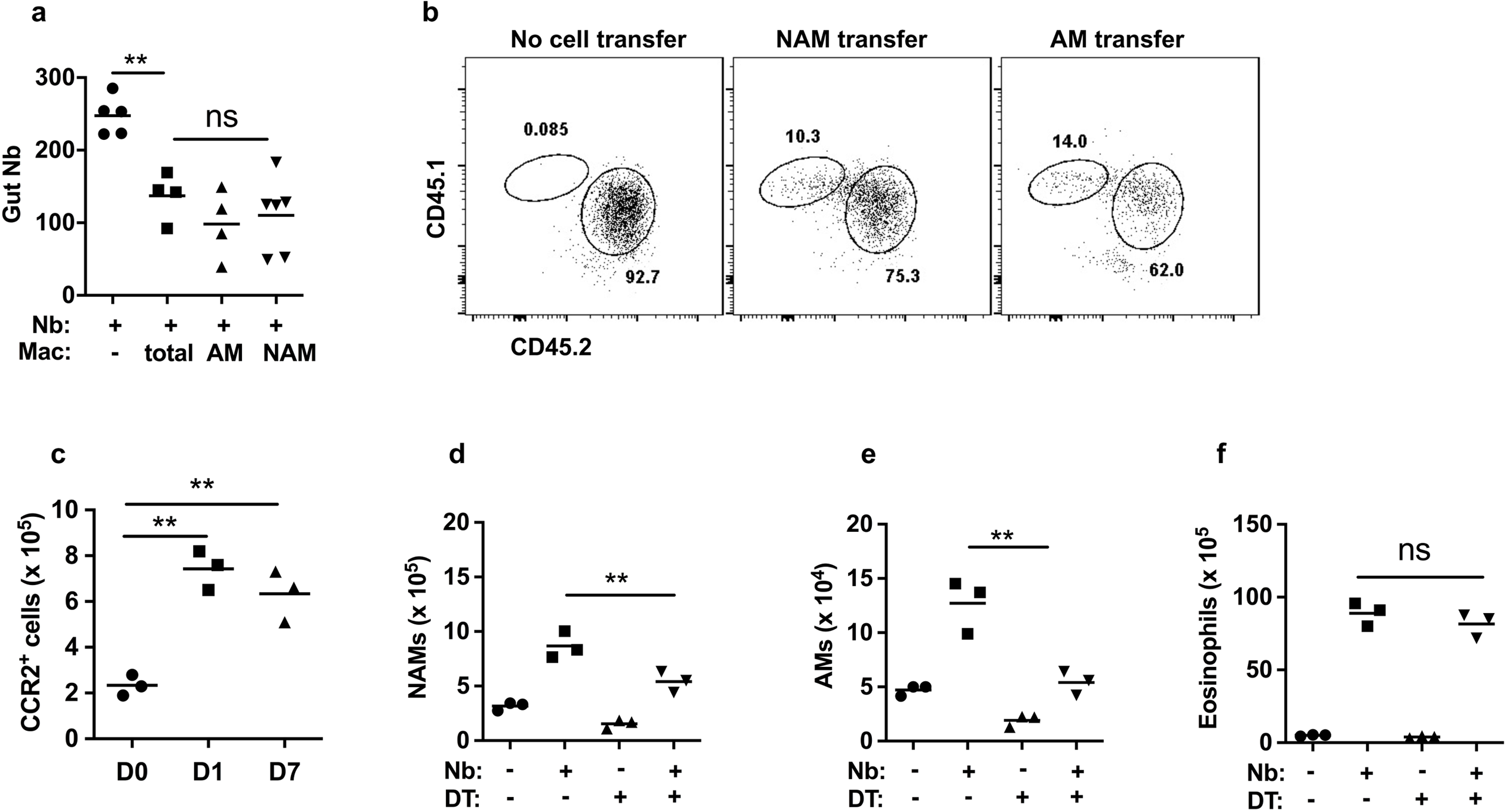
Lung infiltrating monocytes acquire a tissue-resident phenotype and contribute to the increased alveolar macrophage population after *N. brasiliensis* inoculation. (a) At day 7 after *N. brasiliensis* (Nb) inoculation, donor macrophages including total macrophages, alveolar macrophages (AMs), and non-alveolar macrophages (NAMs) were electronically sorted and transferred to recipient mice, which were inoculated with Nb two days later. Recipient mice were assayed for intestinal worm number at day 5 after inoculation; a control group did not receive macrophages. (b) donor CD45.1 mice were inoculated with Nb and 7 days later, lung NAMs and AMs (as described in Fig. 1) were sort-purified and transferred i.t. into naive recipients (CD45.2), which were inoculated with Nb 2 days later. Donor and recipient lung AM populations were assessed at day 5 after Nb inoculation. Flow cytometry plots were representative of 3 mice per group and 2 independent experiments. (c) CCR2-GFP reporter mice were assessed by flow cytometric analysis for monocyte recruitment to total lungs at days 1 and 7 after Nb inoculation. (d-f) Monocytes were depleted by administration of DT at -1, +1, and +3 after Nb inoculation of CCR2-DTR mice. At day 5 after Nb inoculation, numbers of lung NAM (d), AM (e), and eosinophils (f) were determined by flow cytometric analysis. Each symbol represents an individual mouse and horizontal lines indicate the mean. All results are representative of two independent experiments. **p<0.01

To further explore the characteristics and functional significance of monocyte-derived alveolar macrophages (Mo-AMs), we employed established fate mapper mice that allow monocyte-derived cells in adult mice to be identified by their history of *Cx3cr1* expression^24^. With this mouse model, monocytes can be tracked in *Cx3cr1*^CreERT2-IRES-YFP^ mice (abbreviated as *Cx3cr1*^Cre^ mice), which express a tamoxifen (TAM)-inducible Cre recombinase (CreERT2) under the control of the endogenous promoter, followed by an IRES–EYFP element^24, 25^.

To track the fate of the CX3CR1^+^ monocytes, *Cx3cr1*^Cre^ mice were crossed with *Rosa26*stop^-tdTomato^ reporter mice (abbreviated as *R26*tdTomato mice), as previously described^24^. The resultant *Cx3cr1*^CreERT2-IRES-YFP/+^*Rosa26*^floxed-tdTomato/+^ mice were inoculated with Nb and administered TAM at day -1 and day +1 post-infection, which irreversibly labels CX3CR1^+^ cells and their progeny by inducing the expression of tdTomato. As a control, naïve *Cx3cr1*^CreERT2-IRES-YFP/+^*Rosa26*^floxed-tdTomato/+^ mice were also administered TAM (**Fig. 4a**). At day 7 after inoculation, lung cell suspensions were collected and stained for markers associated with conventional AMs (CD64^+^,F4/80^+^,SiglecF^+^,CD11c^+^) (**Fig. 4b - 4c**). The gated AM subset was then analyzed for tdTomato expression. As shown in **Fig. 4d**, ∼30% of AMs were tdTomato^+^ post-Nb infection, indicating that these cells are derived from CX3CR1^+^ monocytes, while AMs from naïve controls similarly treated with tamoxifen showed less than 1% tdTomato expression (**Supplementary Fig. 1a,b**). Interestingly, tdTomato^+^ positive cells persisted for up to a month post-infection (**Fig. 4e**) and appeared phenotypically distinct from tdTomato negative AMs by ultrastructural analysis (**Fig. 4 f**). Importantly, the use of these fate mapping mice obviated the need for cell transfers and/or generation of chimeras for tracing development of Mo-AMs, providing a useful more physiological method for tracking and isolating Mo-AMs and tissue-derived (TD) AMs. Next, we sort-purified TD-AMs (tdTomato^-^) and Mo-AMs (tdTomato^+^) at day 7 after Nb inoculation and cultured them with Nb L3 to test whether they possessed common or unique anti-helminth effector activity. Importantly, tdTomato^+^ AMs derived from CX3CR1^+^ monocytes were significantly more effective at impairing L3 metabolic activity (quantified by decreased ATP) and killing Nb larvae than their tissue-derived counterparts (**Fig. 4g,h**). These studies thus indicate that after Nb inoculation, monocytes enter the lung and transition to an AM-like phenotype by acquiring expression of Siglec-F and CD11c. However, these data also suggest that monocyte-derived AM entering the lung post-infection are functionally distinct cells that preferentially damage and kill invading helminth larvae.

**Fig. 4.**
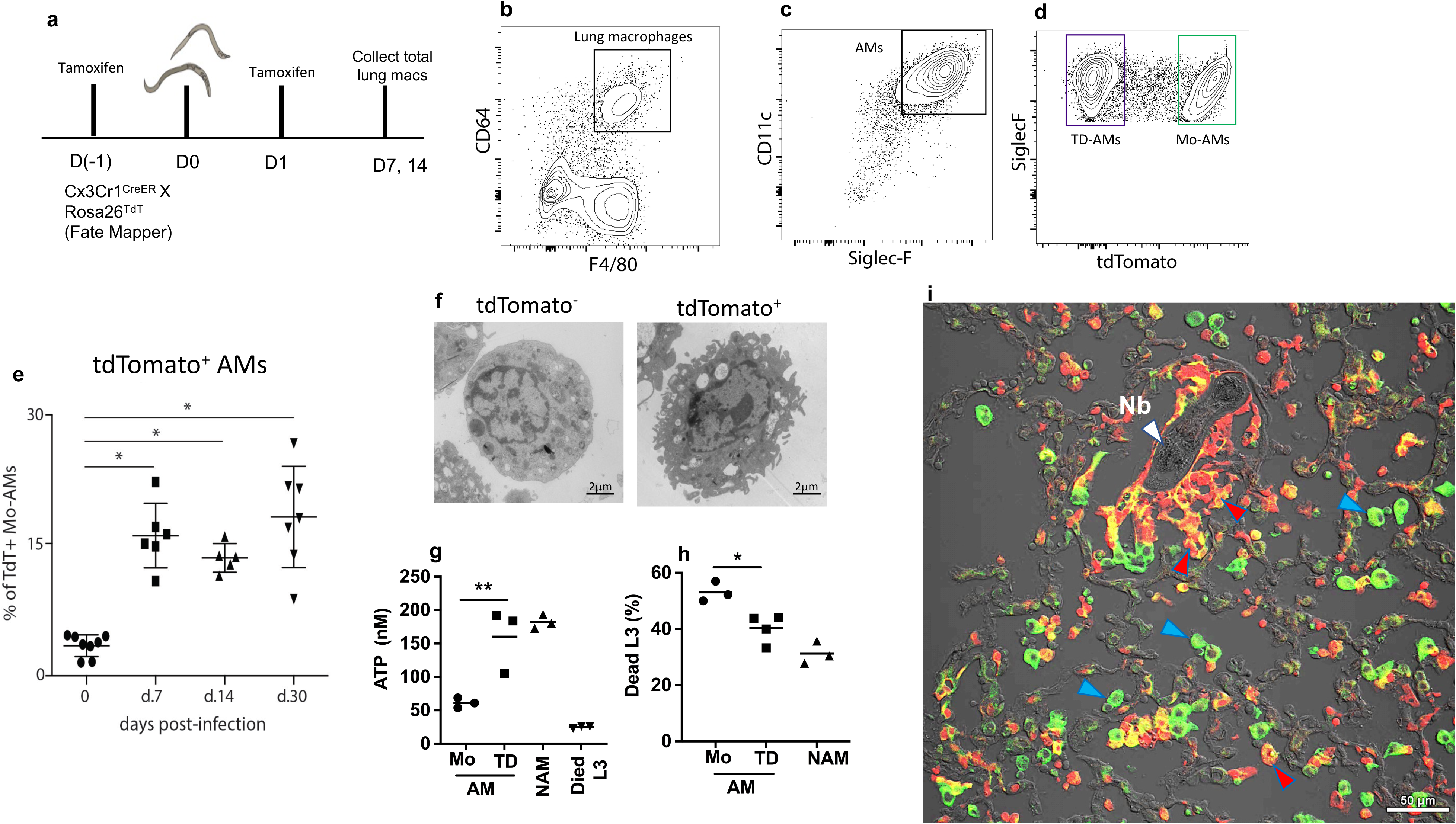
Monocyte-derived alveolar macrophages preferentially damage and kill *N. brasiliensis* L3. (a-d) Fate-mapping of monocyte-derived macrophages recruited to the lung after *N. brasiliensis* (Nb) infection. *Five* mice were pulse administered tamoxifen at days -1 and +1 after Nb inoculation (Fig 1a). Whole lung was collected at day 7 after inoculation and single cell suspensions stained for alveolar macrophages (AMs) as described in Fig. 1. F480^+^,CD64^+^ lung macrophages were gated for expression of CD11c and Siglec-F (b,c) and double positive cells were in turn gated for tdTomato expression (d). (e) Whole lung cell suspensions from *Cx3*cr1^CreERT2-EYFP-IRES-YFP/+^Rosa26^floxed-tdTomato/+^ reporter mice were assessed for tdTomato (tdT)^+^ monocyte-derived alveolar macrophages (Mo-AMs) at different timepoints after Nb inoculation. (f) tdTomato^-^ and tdTomato^+^ cells were sort- purified on day 30 post-infection and prepared for ultrastructural analysis via TEM (magnification 3800x) (g, h) Lung tdTomato ^+^ (tdT^+^ AM) monocyte-derived AMs and tdTomato^-^ (tdT-AM) tissue-derived AMs were sort purified from Nb-primed *Cx3*cr1^CreERT2- EYFP-IRES-YFP/+^Rosa26^floxed-tdTomato/+^ reporter mice at day 7 post-infection. Macrophage subsets were seeded to 24-well plates (1 x 10^6^ cell/well) and co-cultured with 100 exsheathed L3. At day 5 after culture, worm ATP concentration (g) and percent mortality (h) were determined. Triplicate samples were assayed from pools of 20 larva from each well for determination of ATP concentration. Each symbol represents mean of triplicate samples and this experiment was repeated three times with similar results. (i) Lungs from *Cx3*cr1^CreERT2-EYFP-IRES-YFP/+^Rosa26^floxed-tdTomato/+^ reporter mice were collected after a second inoculation with Nb., sectioned, and stained with APC-CD11c antibody for confocal Immunofluorescence imaging analysis of CD11c and CX3CR1 (tdTomato^+^) expression. Image is representative of total 8 individual mice. White marker, Nb larvae; blue marker, CD11c^+^, CX3CR1^-^ TD-AMs, and red marker, CD11c^+^, CX3CR1^+^ Mo-AMs.

To examine the localization of Mo-AMs in lung tissue during Nb infection, *Cx3cr1*^CreERT2-IRES-YFP/+^*Rosa26*^floxed-tdTomato/+^mice were administered TAM as described above and inoculated with Nb or left untreated. Bronchioalveolar lavage (BAL) was collected at day 7 after Nb inoculation. As shown in **Supplementary Fig. 1c**, Mo-AMs were not detected in BAL of uninfected mice, but were readily detectable in the BAL after Nb inoculation. In contrast, TD-AMs were readily detectable in BAL in untreated and Nb inoculated mice, with little change after infection (**Supplementary Fig. 1d**). Remaining lung tissue after collection of BAL was also assessed and although Mo-AMs were not observed in uninfected mice as expected, Mo-AMs and TD-AMs were readily detectable after infection (**Supplementary Fig. 1c,d**). The presence of both AM subsets in remaining lung tissue may be in part due to residual alveolar macrophages in alveoli and airways after lavage as well as potential disruption of lung architecture resulting from infection. The in situ distribution of Mo-AMs was also assessed after secondary inoculation. As shown in **Supplementary Fig.1e**, marked increases in Mo-AMs in the BAL were detected after secondary inoculation, compared to primary inoculation. Collectively, these data suggest that Mo-AMs have populated the lung and are ideally positioned to encounter invading parasitic larvae following a secondary challenge.

To visualize potential interactions between macrophages and larvae, immunofluorescent staining of cryosections from Cx3cr1^CreERT2-IRES-YFP/+^*Rosa26*^floxed-tdTomato/+^mice was assessed after Nb secondary inoculation. In these experiments, the second inoculation was administered 2 days after the first inoculation and mice were sacrificed 6 days after primary inoculation. This approach optimized visualization of trafficking parasites, as parasites are rapidly killed if the second inoculation is given at later time points and are thus not readily visualized. As shown in **Fig. 4i**, using confocal microscopy, Mo-AMs (tdTomato^+^, CD11c^+^; yellow-orange) are observed immediately surrounding the parasite after a second infection, whereas TD-AMs (tdTomato^-^,CD11c^+^;green) are more restricted to the airways. tdTomato^+^, CD11c^+^ cells (Mo-AMs) are also found in the airways consistent with our flow cytometric analysis of BAL fluid. It should be noted that >98% of tdTomato^+^, CD11c^+^ cells exhibited an AM-like phenotype (F480^+^, CD64^+^, CD11c^+^, Siglec-F^+^) as determined by flow cytometric analysis (**Supplementary Fig. 1 f,g**).

Our findings that these Mo-AMs exhibit enhanced anti-helminth activity provokes the hypothesis that they are phenotypically distinct from their tissue-derived counterparts. To further test this, tdTomato^+^ Mo-AMs and tdTomato^-^ TD-AMs were administered TAM (as in **Fig. 4a**), sort-purified on day 7 after Nb inoculation and subjected to global transcriptome profiling. RNA-seq analysis showed that Mo-AMs had a gene expression profile distinct from TD-AMs (**Fig. 5a**). Even by day 14 post-infection Mo-AMs maintained distinct profiles relative to TDAMs and naïve AMs, as determined by pairwise Euclidean distance calculation (**Fig. 5a**) and principal components analysis (**Fig. 5b**). Visualization of overall expression patterns by volcano plots showed more transcriptionally regulated genes in Mo-AMs compared to TD-AMs at day 7 and day 14 (**Fig. 5c**). Venn diagrams of upregulated (**Supplementary Fig. 2a**) and downregulated (**Supplementary Fig. 2b**) genes further illustrated the distinct differences in expressed genes between Mo-AMs and TD-AMs after Nb inoculation. Ingenuity Pathway Analysis (IPA) further suggested that the TD- and Mo-AMs were functionally distinct (**Supplementary Fig. 2c**). Importantly, canonical type 2 pathways that are critical in mediating helminth resistance were more pronounced in Mo-AMs relative to TD-AMs (**Fig. 5d**). Specifically, Arg 1, Retnla, Il-13, and Mgl2 were all increased in Mo-AMs relative to TD-AMs on day 7 and to a lesser extent on day 14 post-infection (**Fig. 5d**). While Mo-AMs remained heightened in these responses on day 14 post-infection, TD-AMs had returned to baseline levels or in some cases below baseline levels by this timepoint (**Fig 5d**). These data suggest that Mo-AMs develop a potent M2-associated phenotype with anti-helminth functions. Additionally, wound healing markers showed a similar pattern of increased expression in Mo-AMs (**Supplementary Fig. 2d**). Notably, peroxisome proliferator-activated receptor gamma (PPARγ) associated signaling pathways, important in lipid metabolism and surfactant homeostasis^26^, were instead significantly reduced in Mo-AMs relative to naïve TD-AMs at day 7 (log_2_ fold change -1.72; FDR p<0.001) and day 14 (log_2_ fold change -1.95; FDR p<0.001)) after Nb inoculation. This may reflect a more prominent role for TD-AMs in homeostatic function associated with lipid metabolism.

**Fig. 5.**
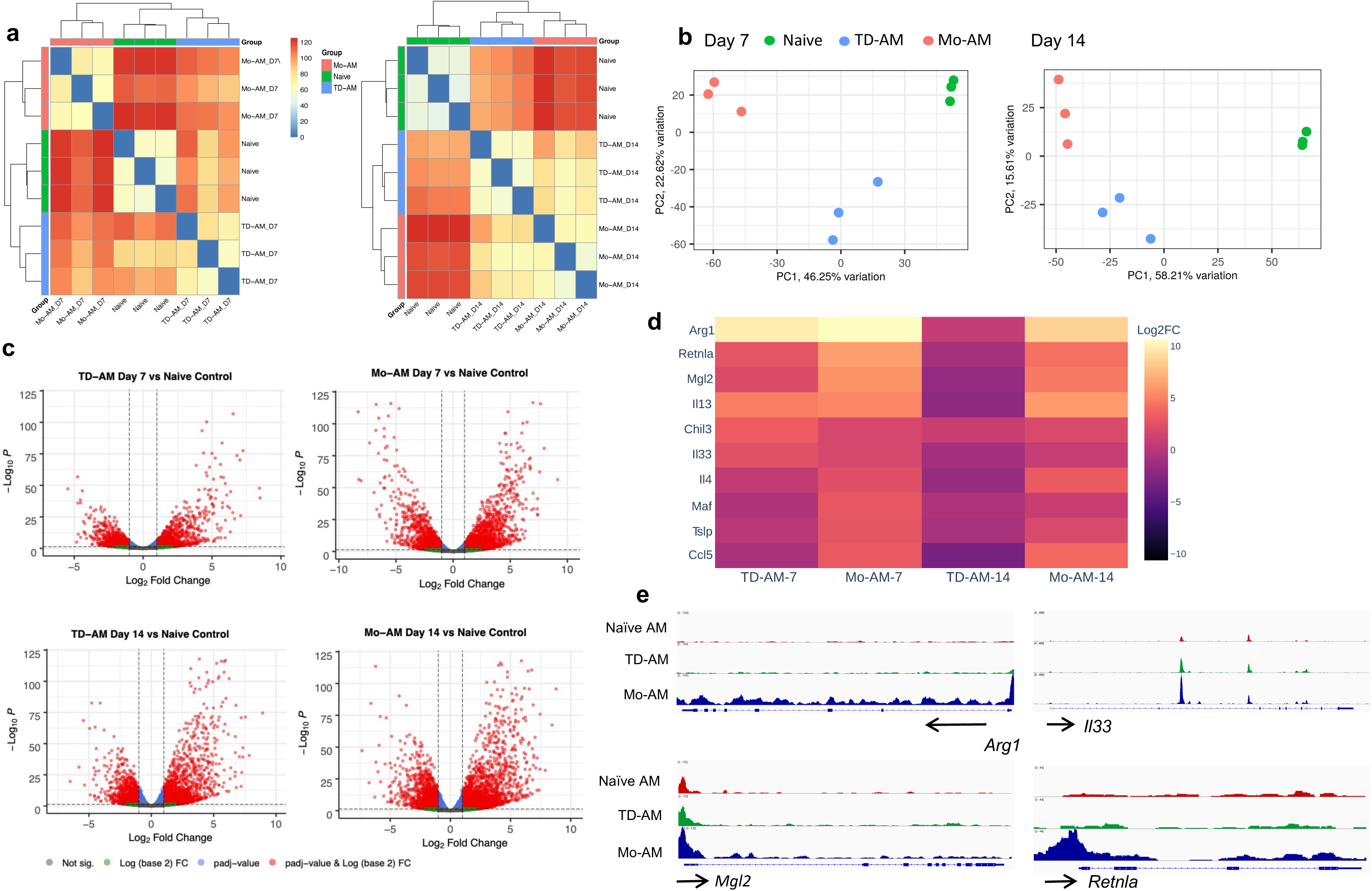
Monocyte-derived alveolar macrophages have distinct phenotype characterized by upregulation of type 2 markers. *Cx3*cr1^CreERT2-EYFP-IRES-YFP/+^Rosa26^floxed-tdTomato/+^ reporter mice received tamoxifen at days -1, and +1 after *N. brasiliensis* (Nb) infection. At day 7 and day 14, tdTomato^+^ monocyte-derived alveolar macrophages (Mo-AMs) and tdTomato^-^ tissue-derived AMs (TD-AMs) were sorted-purified for RNA-seq transcriptional analysis and compared to naïve AMs, with 3 mice/treatment group. (a) pairwise Euclidean distance relative to the transcriptional profiles, demonstrating sample relatedness between AMs from untreated mice, and Mo-AMs and TD-AMs at day 7 and 14 after Nb inoculation (b) Principal Component Analysis of transcriptional profiles of treatment groups; (c) volcano plots of TD-AMs and Mo-AMs individual gene expression profiles at day 7 and day 14 expressed as compared to Ams from naïve mice; (d) Expression of selected characteristic type 2 response markers in TD-AMs and Mo-AMs at day 7 and day 14 after Nb inoculation as expressed relative to AMs from naïve mice(Log_2_ fold change); (e) Genome browser views of the *Arg1*, *Il33*, *Mgl2* and *Retnla* in naïve, TD-AMs, and Mo-AMs. Each track represents the normalized read counts of accessible chromatin regions.

In independent experiments, we also employed transposase-accessible chromatin with sequencing (ATAC-seq) to examine changes in chromatin profiles by identifying transposase-accessible regulatory gene elements in both TD- and Mo-AMs subsets on day 7 post-infection. At a global level, the chromatin profiles of TD-AM from Nb inoculated mice were more similar to naïve alveolar macrophages than Mo-AMs from Nb inoculated mice as indicated by Euclidean distance calculation (**Supplementary Fig. 3a**) and principal component analyses (**Supplementary Fig. 3b**). Differential volcano plot analyses revealed greater accessibility of regulatory elements in the Mo-AMs from Nb inoculated mice (**Supplementary Fig. 3c**), consistent with their increased overall transcription levels as observed with RNAseq (**Fig. 5a-d**). Comparison of differentially called peaks in the four replicates of Mo-AMs versus TD-AMs identified 33,229 differentially expressed peaks (**Supplementary Fig 3d**). No significant differences were seen in chromatin accessibility between *Il33*. However, Mo-AMs had significant increases (p<0.001) in chromatin accessibility at the *Arg1, Mgl2,* and *Retnla* promoter loci **(Fig 5e)**. These findings suggest that the regulation of differential gene expression in TD-AMs is mediated at the level of chromatin accessibility. These data correlate with the increased *Arg1*, *Mgl2*, and *Retnla* transcripts also observed in the Mo-AMs (**Fig. 5d**).

Analyses of in vivo proliferation of macrophage subsets showed similar levels of proliferation between TD-AMs and Mo-AMs at day 7 and day 14 after inoculation (**Supplementary Fig. 4a,b**), consistent with both populations undergoing expansion in part through cell cycling. However, Seahorse real-time cell metabolic analyses using the mitochondria stress test revealed a higher oxygen consumption rate (OCR) and a higher extracellular acidification rate (ECAR), in Mo-AMs compared to TD-AMs at day 7 after inoculation (**Supplementary Fig. 4c**), indicating increased oxidative phosphorylation. We also observed increased *Il13* mRNA in both alveolar macrophage subsets after Nb inoculation. To corroborate these findings, we assessed IL-13 protein cytoplasmic staining and observed significant increases in IL-13 in alveolar macrophages from Nb but not *IL-4Ra*^-/-^ mice at day 7 after inoculation (**Supplementary Fig. 4d**). These data are consistent with previous reports of lung macrophages producing IL-13^27^.

Collectively, our discovery-based studies revealed preferentially elevated levels of *Arg1* gene expression in Mo-AMs, which were also most effective at killing L3 (**Fig 5d**). To corroborate these findings, we performed qPCR on tdTomato^+^ Mo-AMs and tdTomato^-^ TD-AMs obtained from total lungs of *Cx3*cr1^CreERT2-IRES-YFP/+^Rosa26^floxed-tdTomato/+^ mice 8 days after L3 inoculation. As shown in **Fig. 6a**, Arg1 mRNA was markedly elevated in tdTomato^+^ Mo-AMs relative to controls. To assess whether Arg1 protein was similarly increased in this subpopulation, Mo-AMs and TD-AMs were assayed for intracellular Arg1 protein expression by flow cytometric analysis. As shown in **Fig. 6b,c**, Arg1 protein was expressed at considerably higher levels in FACS gated tdTomato^+^ Mo-AMs relative to tdTomato^-^ TD-AMs We further corroborated these findings through cytology and immunofluorescent staining of alveolar macrophage subsets that showed preferential staining for Arg1 in Mo-AMs (**Supplementary Fig. 5**).

**Fig. 6.**
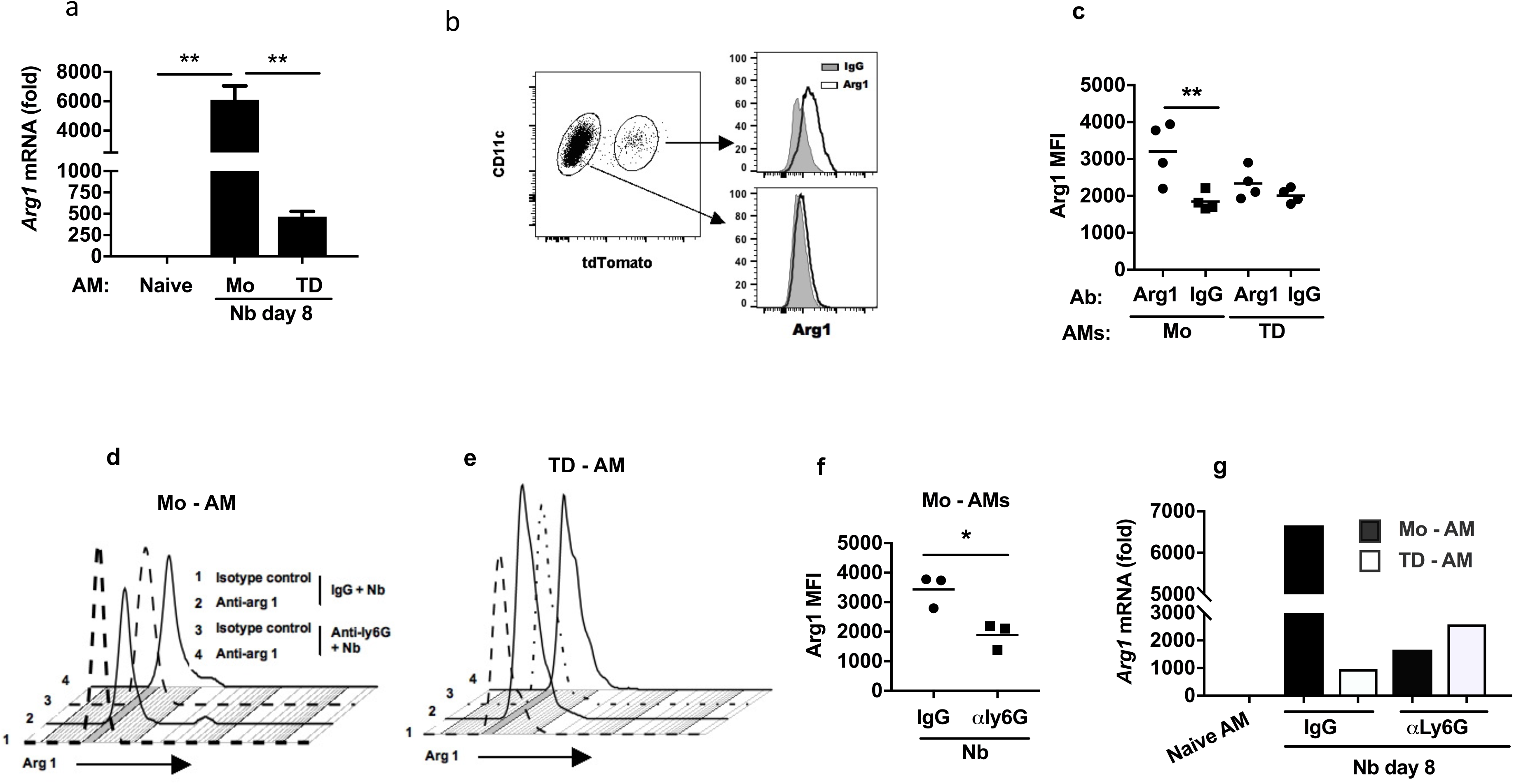
After *N. brasiliensis* infection, monocyte-derived alveolar macrophages preferentially express arginase 1 which is dependent on extrinsic neutrophil signaling. (a-c) *Cx3*cr1^CreERT2-EYFP-IRES-YFP/+^Rosa26^floxed-tdTomato/+^ reporter mice were administered tamoxifen and inoculated with *N. brasiliensis* (Nb) as described in Fig. 5. (a) Arginase 1 (Arg1) mRNA levels by qPCR in subpopulations of lung alveolar macrophage (AM) subsets at day 8 after Nb inoculation expressed relative to naïve AMs; (b) Representative flow cytometric analysis of Arg1 intracellular protein expression in AM subsets at day 7 after Nb inoculation and (c) Arg 1 mean fluorescence intensity (MFI) in individual samples from different treatment groups; (d-g) *Cx3*cr1^CreERT2-EYFP-IRES-YFP/+^Rosa26^floxed-tdTomato/+^ reporter mice were administered anti-Ly6G Ab or isotype control at days -1,1,and 3 and given tamoxifen at days -1 and 1 after Nb inoculation. (d-f) representative flow cytometric analyses of Arg1 intracellular protein expression after neutrophil depletion (f) Arg1 MFI in individual samples from each treatment group. (g) Arg1 mRNA levels by qPCR in macrophage subsets, presented as the fold increase over naïve AMs. Data shown are the mean and SEM of triplicate samples (a) or of duplicated samples (g) from a pool of four to eight mice per group and are representative of two independent experiments. Each symbol represents an individual mouse and horizontal lines indicate the mean. All results are representative of two independent experiments. *p<0.05, **p<0.01

M2 macrophage activation is regulated by several factors including the production of type 2 cytokines by specialized innate immune cells. More specifically, our work and that of others have shown that neutrophils can contribute to M2 cell activation^3, 7^. Therefore, we sought to assess whether the high Arg1 expression was also dependent on Nb-activated neutrophils. A role for neutrophils in preferential activation of Mo-AMs would be consistent with a role for extrinsic signals in addition to intrinsic cell lineage differences contributing to differential macrophage subset activation in the lung. To test this possibility, *Cx3cr1*^CreERT2-IRES-YFP/+^*Rosa26*^floxed-tdTomato/+^mice were administered TAM as described above and anti-Ly6g neutrophil depleting Ab or isotype control was administered at days -1, +1, and day 3 after Nb inoculation. At day 5 after inoculation total lung cell suspensions were stained for AMs and analyzed by flow cytometry for tdTomato expression. Elevations in intracellular Arg1 protein expression (**Fig. 6d-f**) and gene expression (**Fig. 6g**), as measured by RT-PCR, were significantly reduced in tdTomato^+^ Mo-AMs after neutrophil depletion.

Previous studies have shown that macrophage mediated helminth killing is dependent on Arg enzymatic activity through use of specific inhibitors^3, 8^. We next examined macrophages from Arg1 conditional knockout mice, where macrophages were isolated from Tie2-Cre Arg1^fl/fl^ or control Tie2-Cre mice at day 7 after inoculation and cultured with Nb L3. Use of Tie-2 to drive macrophage deletion of *Arg1* has significant advantages over LysMCre, which is ineffective at deleting *Arg1* in macrophages^28^. As shown in **Figure 7a,b**, macrophages from Nb primed Tie2-Cre *Arg1*^fl/fl^ mice showed markedly impaired L3 killing and corresponding increases in metabolic activity as assessed by ATP worm levels. This is consistent with previous studies showing reduced impairment of *Heligmosomoides polygyrus* larval mobility by macrophages from Tie2-CreArg1^fl/fl^ mice^11^. However, it remained unknown how Arg1 actually affects parasitic nematode larval survival. Arg1 triggers a cascade of effects resulting in production of potentially toxic urea and also polyamines, which can promote cellular activation and proliferation^20^. Recent studies of the intestinal and hepatic type 2 responses in Schistosome-infected mice have further shown an important role for Arg1 in depleting local arginine concentrations, and thereby attenuating activation of nearby immune cells, effectively resulting in immunosuppression^21^. Arginine depletion and not metabolite generation mediated the downregulatory effect^29^. Based on these findings, we hypothesized that one mechanism through which Arg1 may kill L3 would be through localized depletion of arginine, an essential amino acid for nematodes, as they are generally not thought to be capable of arginine biosynthesis^30, 31^. To test whether L-arginine was depleted in the macrophage/L3 cocultures L-arginine was directly measured following culture of L3 alone, or coculture of exsheathed L3 with either naïve lung macrophages or lung macrophages from mice inoculated with Nb for 7 days. Significant decreases in L-arginine were detected only in L3 cocultures with lung macrophages from Nb inoculated mice (**Fig. 7c**). We next performed L-arginine add back experiments to examine whether macrophage killing could be blocked by supplementing the media with highly purified L-arginine. As shown in **Fig. 7d, e**, larval killing was significantly decreased with increased L-arginine supplementation and live parasite metabolic activity, as measured by ATP values, was markedly increased. To assess whether L-arginine was required for L3 survival under in vitro conditions, exsheathed larvae were cultured in arginine-free media. As shown in **Fig 7f**, parasite mortality was high in arginine-free media, while addition of arginine significantly reduced mortality. Furthermore, supplementation of arginine free media with exogenous L-arginine enhanced parasite ATP levels (**Fig. 7g**). To examine whether anti-helminth functions of lung macrophages were specific to Nb, primed lung macrophages isolated at day 7 after Nb inoculation were cocultured with a different nematode parasite, *H. polygyrus*. L3 mortality (**Fig. 7h**) was markedly increased and was reduced by supplementation with L-arginine. Similarly reduced L3 ATP levels were markedly enhanced by L-arginine administration (**Fig. 7i**). These studies thus indicate that Nb-primed macrophages are capable of killing other nematode parasite species and provide substantial insight into the protective mechanisms of M2 macrophages.

**Fig. 7.**
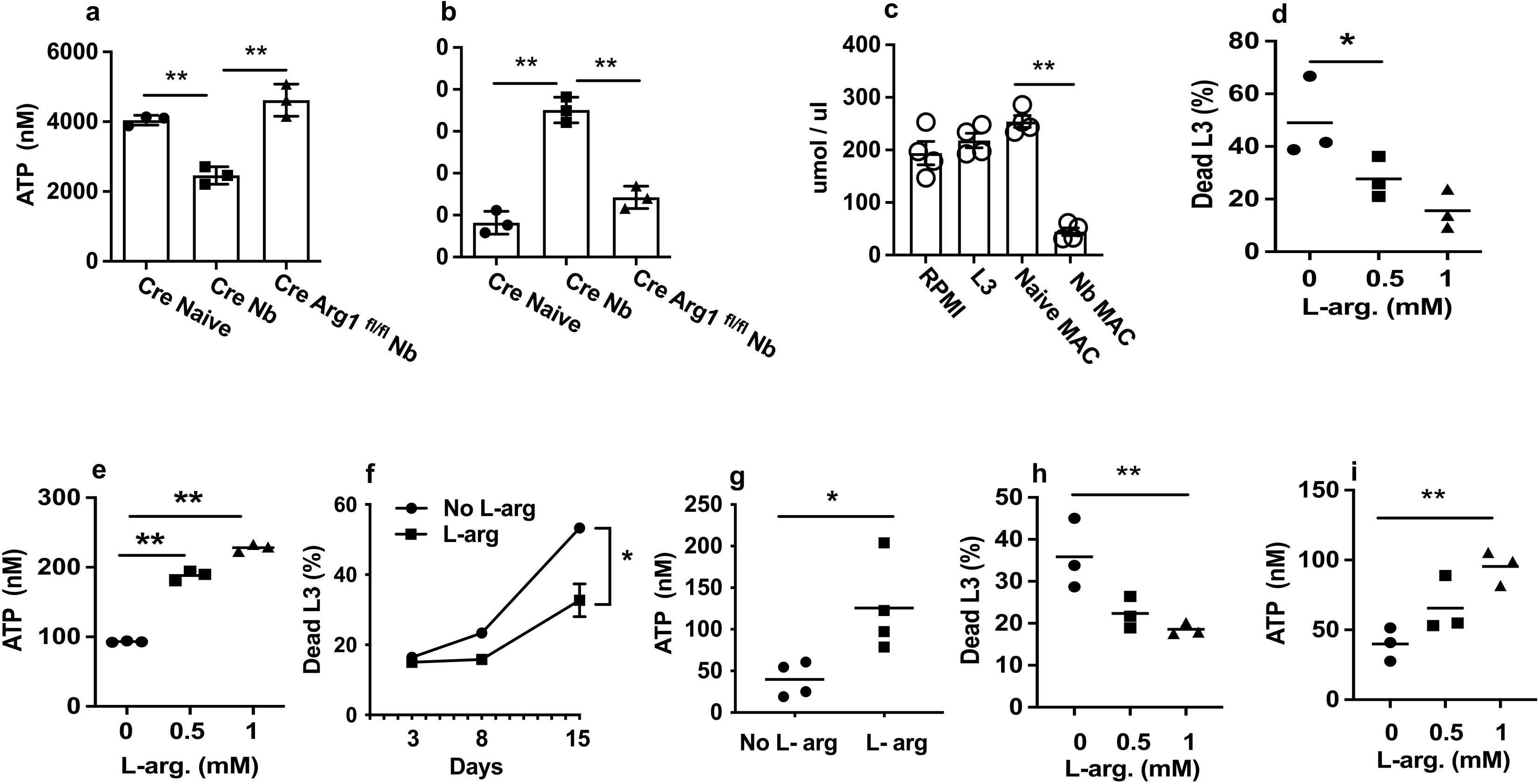
L-arginine depletion by Nb activated lung macrophages kills parasitic larvae. (a,b) Tie2-Cre (CRE) and Tie2-Cre Arg1^fl/fl^ (Cre Arg1^fl/fl^) mice were inoculated with Nb L3 and 5 days later total lung macrophages were sort purified and cultured (1X10^6^ cells/well) with 100 Nb exsheathed L3 and assayed for worm ATP (a) and percent mortality (b). (c) Arginine levels in supernatant were measured in cultures with L3, or cocultures with sort-purified naïve lung macrophages or lung macrophages isolated from Nb, at day 5 after culture. (d,e) Sort purified total lung macrophages from Nb inoculated mice were cocultured with Nb exsheathed L3 naïve macrophages and supplemented with L-arginine. At day 5 after culture, the percent mortality (**d**) and larval ATP concentration (e) were determined, as described in Fig. 4. (f, g) ex-sheathed Nb larva (100 L3) were seeded to 12 well plates and cultured with L-arginine free RPMI1640 media, with one group supplemented with L-arginine. At day 3, 8, and 15 after culture, the percent mortality was assessed (**f**), and remaining worm ATP concentration assessed at day 15 (**g**). Data are presented as the mean and SEM of triplicate samples (f) or presented as individual samples obtained from 20 larva from each well for determination of ATP concentration and horizontal lines indicate the mean(g). (h,i) Total lung macrophages were sort-purified from Nb inoculated mice as described above and co-cultured with L3. Percent mortality (h) and worm ATP concentration (i) were assessed. In all experiments, each symbol represents an individual mouse and horizontal lines indicate the mean. All results are representative of ate least two independent experiments. *p<0.05, **p<0.01

## Discussion

Macrophages mediate resistance to helminths as well as microbial pathogens. Recent studies suggest that the lung is an important target for therapeutic strategies and vaccine development against helminths^2^, and that lung macrophages with anti-helminth effector functions develop shortly after Nb infection^3^. Tracking monocyte recruitment to the lungs and their differentiation in this microenvironment, we showed that during infection they can develop an alveolar macrophage-like phenotype that is distinct from TD-AMs and that preferentially mediates helminth killing and includes expression of high levels of Arg1, which is critical for helminth resistance. Development of this polarized Mo-AM phenotype was dependent on neutrophil help and our studies further show that macrophage-derived Arg1 mediates parasite killing through localized depletion of arginine, providing a previously unappreciated nutrient deprivation mechanism for macrophage-mediated helminth killing.

Emerging studies have now demonstrated that macrophages exhibit tissue-specific phenotypes that are ideally suited for their specific microenvironments^18, 24, 32, 33^. In the lung, TD-AMs play a critical role in lung homeostasis including clearance of debris and surfactant recycling^16, 34^. After Nb infection, the lung NAMs are a heterogeneous population, with macrophages derived and from monocytes and also tissue-derived interstitial macrophage populations ^17^. Our findings with the CCR2-DTR mice, and also with the transfer experiments using CD45 congenic mice, now indicate that the rapid increase in alveolar macrophages observed after Nb infection is supported by monocyte-derived cells. Furthermore, these Mo-AMs are differentially activated with distinct effector functions including enhanced parasite killing and Arg1 expression, thereby explaining why NAMs were less effective at killing the helminths in vitro, but after transfer, in the context of the lung microenvironment, could develop this anti-helminth effector function associated with their expression of AM surface markers. Recent studies in a chimeric model showed that recruited Mo-AMs during influenza infection preferentially conferred protection against heterologous bacterial infections^18^. Interestingly, fibrosis was also linked to a recruited Mo-AM population with increased Arg1 expression in an acute lung injury model ^19^. Our studies now show that tracking of monocytes, using fate mapping mice under physiological conditions, reveals their differentiation into Mo-AMs that are characterized by high Arg1 expression and enhanced anti-helminth effector function. Parasites typically leave the lung and migrate to the small intestine by day 4 after inoculation^35^. The magnitude of the type 2 inflammatory response decreases shortly thereafter, but still remains potent, resulting in chronic inflammation and the associated development of fibrosis and emphysematous pathology^36, 37^. Mo-AMs expressed high levels of CD11c and Siglec-F by day 7 but transcriptome analyses revealed that they were distinct from TD-AMs as revealed by PCA and Euclidean distance analyses. The observation that this population expressed considerably higher levels of Arg1 and other type 2 markers compared to TD-AMs indicates a distinct activation state and effector function. By day 14, upregulated gene expression of immune response genes was decreased in the TD-AMs, while Mo-AMs retained relatively high expression of these genes that including type 2-associated molecules. These findings are consistent with a model where TD-AMs revert back to a housekeeping phenotype shortly after infection, likely to assume their critical function of maintaining homeostasis and healthy lung function^16^. In this context, as described in the Results, PPARγ expression remained higher in TD-AMs at both day 7 and day 14 after Nb inoculation relative to Mo-AMs. PPARγ mediates lipid metabolism, a critical homeostatic function of TD-AMs^16, 26^. However, our findings may also be consistent with recent findings that TD-AMs may undergo paralysis associated with poor phagocytic activity after bacterial infections^38^. Further studies of TD-AM function after helminth infection are needed to distinguish between these possibilities. In contrast AMs derived from monocytes maintain a more activated phenotype with high expression of M2 markers and associated remodeling of the chromatin landscape. As such the Mo-AMs are likely primed to respond to subsequent helminth infections. In future studies it will be interesting to examine the duration of changes in the chromatin landscape relative to gene expression, as these epigenetic modifications may potentially contribute to trained immunity, consistent with previous studies showing persistence of trained macrophages as late as 45 days after Nb inoculation^3^. These epigenetic changes in macrophages may also affect heterologous infections, consistent with previous studies that Nb infection can enhance susceptibility to infection with Mycobacterium tuberculosis in part through impaired macrophage effector function^39^. Interestingly, we also observed increases in macrophage expression of IL-13, which was confirmed by cytoplasmic IL-13 staining. Previous studies have also shown increased lung macrophage IL-13 production during type 2 responses, in some cases directly contributing to the polarization of the response^27, 40^. It is increasingly apparent that a number of different innate myeloid and lymphoid immune cells can express type 2 cytokines during allergic and helminth responses, likely triggered by activation of common pathways including IL-4R signaling^10^, and potentially playing essential roles at different stages of the response.

After primary infection, our findings show that several days elapse as the monocytes are recruited to the lung and differentiate to the alveolar macrophage phenotype. The Mo-AM population likely has little effect on parasite migration during a primary infection, consistent with most parasites successfully reaching the small intestine by day 4 after primary inoculation. However, after infection, the Mo-AMs persist in the lung for prolonged periods: they are readily detected as late as 30 days after infection (data not shown). As such, during a subsequent infection, parasites can encounter these macrophages in the lung, slowing their migration and eventually culminating in their morbidity and death in the lung over a period of several days, thereby blocking their migration to the intestine. Our findings further suggest that the Mo-AMs play a critical role in contributing to this accelerated resistance, likely through their heightened Arg1 activity, which depletes L-arginine levels. We now show that these Mo-AMs are in close proximity to Nb larvae in the lung after infection. Whether the macrophages simply adhere to the parasite as it migrates through the lung, the larvae are trapped in aggregates of macrophages after infection, or the macrophages potentially swarm the parasite is unknown. It should be noted that increased Mo-AM activation including expression of additional genes associated with type 2 immunity may also trigger other effector mechanisms contributing to anti-helminth effects. The overall primed state of Mo-AMs may also play a significant role in infections with heterologous pathogens, potentially providing protective immunity or alternatively increased susceptibility as has been observed with *Mycobacterium tuberculosis* infections^39^. As such, heterogeneity between individuals in innate responses to specific pathogens may be potentially influenced by previous heterologous pathogen exposure. The genetic background of the host may also be of importance and our studies of Mo-AMs have necessarily focused on BL/6 mice as that is the genetic background of the fate mapping mice used in these studies. Previous studies have shown potent type 2 responses to *N. brasiliensis* in both BL/6 and BALB/c mice^3, 6, 36^. In future studies, the role of Mo-AMs in mediating helminth resistance in mice with varying genetic backgrounds should be investigated.

Neutrophils are increasingly recognized as significant players in helminth infections and in cross talk with macrophages. Recent studies indicate that neutrophils exhibit a distinct alternatively activated (N2) phenotype during helminth infections^3^ and that they can contribute resistance mechanisms leading to reduced worm burden after primary inoculation^4, 5^. Interactions of macrophages with neutrophils can promote M2 activation, likely through both production of type 2 cytokines and direct cell-cell interactions involving efferocytosis^3, 7^. Also, in the lung environment, surfactant protein A can also promote M2 macrophage activation^41^. Interestingly antibodies are not required for resistance to Nb^23^, although they do play an important role in resistance and macrophage Arg1 expression during *H. polygyrus* infection^11, 23^. The lung tissue microenvironment, including neutrophil interactions, thus provides sufficient signals for M2 macrophage activation even in the absence of Ab signaling. Effects of neutrophil-macrophage crosstalk in driving activation and associated effector function of these myeloid cells can play a significant role in immune responses against diverse pathogens^42^. Our findings indicate that neutrophils are critical for the polarization of the Mo-AM subset towards elevated expression of type 2 markers, including Arg1. Our findings that neutrophils preferentially stimulate this phenotype in recruited Mo-AMs indicates that both intrinsic lineage-specific activation signatures and extrinsic signals provided in the inflamed lung tissue microenvironment interact to drive their activation culminating in an anti-helminth effector phenotype.

Although previous studies have identified Arg1 as being essential in macrophage-mediated helminth killing in the skin^8^, lung^3^, and small intestine^9, 11^, the mechanism through which this enzyme mediates this effect has remained unknown. Our studies now reveal a mechanism that is independent of the many molecules produced during the Arg1 cascade culminating in production of urea, ornithine and polyamines. Our findings do not exclude other mechanisms involving the Arg1 pathways, or potentially other molecules produced independently of Arg1. However, our studies do indicate that Arg1-mediated l-arginine depletion is essential in macrophage-mediated killing of L3. The production of high levels of Arg1 by macrophages as they surround the invading parasitic larvae deprives the parasite of arginine. We now specifically show that Nb metabolism is slowed, and mortality markedly increased in the absence of an external source of arginine in Nb larval cultures with primed M2 macrophages. The developing L3 likely obtains this essential amino acid from the vertebrate host as it traffics through various tissues in vivo. The ability of the activated M2 macrophage to adhere to the invading L3 results in a localized depletion of this critical nutrient, effectively providing a potent anti-helminth effector mechanism. Our findings that arginine supplementation was sufficient to block macrophage-mediated killing in both in vitro cultures and in vivo indicates the potent effect of this anti-helminth macrophage effector mechanism. Given the intense selection pressure of helminths on vertebrate evolution, one can postulate that the high level of Arg1 produced by M2 macrophages, specifically helminth activated Mo-AMs, may in part be an adaptation resulting in a potent resistance mechanism against these multicellular pathogens. In contrast, Arg1 does not appear to significantly impact lung immune cell activation and inflammation during type 2 responses, including schistosome egg-induced granuloma formation^43^. It is intriguing that in microbial infections arginine also plays a critical role in resistance, through generation of nitric oxide synthase 2 activity resulting in the production of anti-microbial nitric oxide. As such, arginine metabolism thus plays an important role in mediating resistance against eukaryotic as well as microbial pathogens although the mechanisms contributing to the protective response are quite different. The mechanism of Arg1-mediated resistance is indeed more similar to Arg1-mediated suppression of Th2 cell activation, where nutrient depletion also plays a major role.

## Material and methods

### Mice

BALB/c ByJ (CD45.1), BALB/c (CD45.2), *Rosa26*^floxed-tdTomato^ and *Cx3cr1*^CreERT2-IRES-YFP^ BL/6 mice were purchased from The Jackson Laboratory (Bar Harbor, ME) and CCR2-DTR, CCR2-GFP, Tie 2-Cre and Tie2-Cre Arg^fl/fl^ mice were all bred and maintained in a specific pathogen-free, virus Ab-free facility at Rutgers New Jersey Medical School Comparative Medicine Resources. Healthy 8-12 week old mice were selected for treatment groups from purchased or bred colonies, without using specific randomization methods or specific blinding methods. The studies have been reviewed and approved by the Institutional Animal Care and Use Committee at Rutgers-the State University of New Jersey. The experiments herein were conducted according to the principles set forth in the Guide for the Care and Use of Laboratory Animals, Institute of Animal Resources, National Research Council, Department of Health, Education and Welfare (US National Institutes of Health).

### Parasite culture and inoculation of mice

*N. brasiliensis* (Nb) L3 were maintained in a petri dish culture containing charcoal and sphagnum moss. The larvae were isolated from cultures using a modified Baermann apparatus with 400U penicillin, 400 μg ml*^−^*^1^ streptomycin, and 400 μg ml*^−^*^1^ Neomycin (GIBCO, Rockville, MD) in sterile PBS, and then washed with sterile PBS three times. Mice were inoculated subcutaneously with a 40µl suspension of 650 Nb L3. For *H. polygyrus*, after propagation, L3 larvae were maintained in PBS at 4°C.

### Flow cytometry and adoptive transfer

Lung tissue was washed with stirring at room temperature for 10 min in Hank’s balanced salt solution (HBSS) with 1.3 mM EDTA (Invitrogen), then minced and treated at 37°C for 30 min. with collagenase (1 mg / ml; Sigma) in RPMI1640 with 10% fetal calf serum (FCS) and with 100 μg / ml of DNase for 10 min. Cells were lysed with ACK (Lonza, Walkersville, MD) to remove erythrocytes. Cells were blocked with Fc Block (BD Biosciences, San Jose, CA), directly stained with fluorochrome-conjugated antibodies against CD45,F4/80, CD64, CD11c, Siglec-F,CD11b,Ly6C, and analyzed by flow cytometry. For adoptive transfer, macrophages (F4/80^+^CD64^+^SiglecF ^vari^CD11c ^vari^), alveolar macrophages (AMs) (F4/80^+^CD64^+^SiglecF^+^CD11c^hi^), or nonalveolar macrophages (NAMs) (F4/80^+^CD64^+^SiglecF^-^CD11c ^vari^), were electronically sort purified (>98%), 5 million cells were transferred i.t. into recipient mice. Two days after cell transfer, the recipients were inoculated with Nb L3. For *Cx3cr1*^CreERT2-IRES-YFP^ x *Rosa26*^floxed-tdTomato^ reporter mice, tamoxifen (75 mg/kg body weight, Sigma, cat#T5648) were delivered to mice at days -1, and +1 after Nb inoculation. At different timepoints after inoculation, single cells were isolated from whole lung and directly stained with fluorochrome-conjugated antibodies against CD45, F4/80, CD64, CD11c, Siglec-F and analyzed by flow cytometry. Cells were gated on CD45^+^,F4/80^+^,CD64^+^ cells, then further gated on Siglec-F^+^, CD11c^hi^ AMs. AMs were separated into tdTomato^+^ monocyte-derived alveolar macrophages (Mo-AMs) and tdTomato^-^ alveolar macrophages (TD-AMs). For intracellular staining of Arg1, cells were first surface stained and then fixed and permeabilized using a kit from BD BioSciences (cat# 554714). For cytospin analyses, sorted cells (2 x10^5^) from lung tissue were suspended in 200 μl of 1X PBS with 2.5% FCS. The sorted cell suspensions were loaded into a Shandon Cytospin 4 (Thermo Electron Corporation, Waltham, MA), spun at 800-1000 rpm for 5 min and stored at -80°C. Frozen cytospin slides were thawed at room temperature for 30 min, fixed in 4% PFA for 15 min, and stained with an APC-conjugated antibody specific to Arginase 1 or corresponding isotype control (Bioss Inc., Woburn, MA). Coverslips were applied to the slides using Vectashield mounting medium (Vector Laboratories, Burlingame, CA) with DAPI. Images were taken using a Leica DM6000B fluorescent microscope, Orca Flash 4.0 mounted digital camera (Hamamatsu Photonics K.K., Japan) and LAS Advanced Fluorescence software (Leica Microsystems, Buffalo Grove, IL). Fluorescent channels were photographed separately and then merged. Exposure times and fluorescence intensities were normalized to appropriate control images.

### Confocal microscopy

Cx3cr1^CreERT2-IRES-YFP/+^Rosa26^floxed-tdTomato/+^ received tamoxifen at day -1 and +1 after primary Nb inoculation of 500 L3. At day 2 after primary inoculation, mice were given a second inoculation of 250. Lung tissues were then collected at day 6 after primary inoculation. This regimen was found to optimize visualization of migrating parasites interacting with primed macrophages. Lung tissues were fixed in 1% paraformaldehyde overnight at 4°C. Tissues were washed with 5mM NH_4_Cl and incubated in 30% (w/v) sucrose overnight at 4°C. Lung tissue was embedded in Optimal Cutting Temperature Compound (Tissue-Tek) and 5 μm sections were cut on CM1950 Cyrostat (Leica, Weltzlar, Germany). Sections were blocked with 1% Rat serum and 1% FcBlock in PBS for 1 hr, followed by 5ug/ml of anti-CD11c-APC overnight and sealed with ProLong Gold Antifade (Invitrogen, Waltham, MA). Confocal Images were captured using a Nikon A1R SI confocal microscope with the DUG GaAsP detector. 10x Plan Apo Lambda N.A. 0.45, 20x Plan Apo VC N.A. 0.75 and 60x Plan Apo VC N.A. 1.4 objectives were used along with DIC optics for the transmitted light images.

### Electron microscopy

Samples were fixed in 2.5% glutaraldehyde / 4% paraformaldehyde in 0.1M cacodylate buffer and then post-fixed in buffered 1% osmium tetroxide. Samples were subsequently dehydrated in a graded series of acetone and embedded in Embed812 resin. 90nm thin sections were cut on a Leica UC6 ultramicrotome and stained with saturated solution of uranyl acetate and lead citrate. Images were captured with an AMT (Advanced Microscopy Techniques) XR111 digital camera at 80Kv on a Philips CM12 transmission electron microscope.

### Macrophage co-cultures with parasitic L3

Macrophages, alveolar macrophages, or nonalveolar macrophages, were electronically sorted from mice inoculated with Nb for 7 days, and placed on 12-well plates (1 x 10^6^) in 2 ml of RPMI1640 medium with 10% of FBS, 400U penicillin, 400 μg/ml streptomycin, and 100 μg/ ml gentamycin. 100 μl of serum from the donor mice was also added to medium. One hundred ex-sheathed L3 larvae were added to each well. Cells and worms were co-cultured for 5 days at 37°C. The procedure to ex-sheath larvae included incubation of L3 with 6.7 mM sodium hypochlorite in PBS for 15 min. at room temperature, washing the L3 six times with sterile PBS containing 400U penicillin and 400 μg ml−1 streptomycin. To ex-sheath Hp L3, the larvae were maintained in phenol RPMI1640 with 10% FBS at 37^0^C, and CO_2_ was added to the medium until yellow color obtained, and then incubated for 25hrs in CO_2_ incubator. The ex-sheathed larvae were isolated using a modified Baermann apparatus with 400U penicillin, 400 μg ml*^−^*^1^ streptomycin, and 400 μg ml*^−^*^1^ Neomycin (GIBCO, Rockville, MD) in sterile PBS, and then washed with sterile PBS three times. In some experiments, the L3 were cultured in L-arginine free RPMI1640 (Thermofisher) without cells and in some groups commercial L-arginine was added to the medium. The ATP levels for the L3 were performed according to the manufacturer’s instructions. Briefly, 20 Nb L3 were collected with 100 μl of PBS and 100μl of the Celltiter-Glo Luminescent reagent (Promega, Madison, WI) was then added. The larvae were homogenized and incubated for five minutes at room temperature to stabilize the luminescence signal. After the homogenate was centrifuged at 1000g for 2 minutes, 100 μl of supernatant was applied to the luminometer to measure luminescence. As a negative control, larvae in RPMI1640 medium were treated in boiling water for five minutes and after cooling, the worms were homogenized with reagent. Dead larvae after incubation with macrophages were defined as non-motile, outstretched bodies, with non-refractive internal structures. L-arginine was measured using an L-arginine assay kit (Abcam, cat#ab252892),

### Cytokine gene expression by RT-PCR

For qPCR, RNA was extracted from lung tissue or sorted macrophages and reverse transcribed to cDNA. qPCR was done with Taqman (Life Technologies Corporation, Carlsbad, CA) kits and the Applied Biosystems QuantStudio 6 Flex Real-Time PCR System. All data were normalized to 18S ribosomal RNA, and the quantification of differences between treatment groups was calculated according to the manufacturer’s instructions. Gene expression is presented as the fold increase over naïve WT controls.

### RNA sequencing

Alveolar macrophages from *Cx3*cr1^CreERT2-IRES-YFP/+^Rosa26^floxed-tdTomato/+^ Nb-inoculated mice were sort-purified by gating on CD45^+^,CD64^+^,F4/80+,CD11c^+^,Siglec-F^+^ AMs then sorted based on tdTomato expression, with TD-AMs being tdT^-^ and Mo-AMs tdT^+^. Cells from a minimum of three biological replicates were sorted and pooled. The purity of all cell populations was 98% or greater. RNA was extracted using the RNeasy Plus Micro Kit (catalog no. 74034; QIAGEN). Illumina-compatible libraries were generated using the NEBNext Ultra II DNA Library Prep Kit for Illumina (catalog no. E7645S; New England BioLabs) and sequenced using an Illumina NovaSeq 6000 system in a paired-end 2x50- base pair (bp) reads configuration. Bulk RNA-seq analysis was performed in accordance with the nf-core RNA-seq guidelines v.1.4.2^44^. Briefly, the output reads were aligned to the GRCm38 (mm10) genome using STAR, followed by gene count generation using feature Counts and StringTie^45–47^. Read counts were normalized and compared between groups for differential gene expression using DESeq2 with significance cutoff at false discovery rate-adjusted p<0.05^48^. Determination of functional pathways was performed using Ingenuity Pathway Analysis (IPA) on differentially expressed genes.

### Assay for transposase-accessible chromatin with sequencing (ATAC-seq)

Sorted cells were lysed in buffer (10 mM Tris-HCl, 10 mM NaCl, 3 mM MgCl_2_, 0.1% octylphenoxypolyethoxyethanol, 0.1% digitonin and 0.1%IPGAL) for 3 min at 4°C and then washed with lysis buffer without digitonin or IPGAL and centrifuged and resuspended with transposase reaction mix (Ilumina Nextera), and incubated for 30 min at 37°C. Cells were then PCR amplified in KAPA HiFi 2× mix (Kapa Biosystems) with barcoding primers. Amplification was conducted for 45 s at 98°C, followed by five cycles of denaturing at 98°C for 15 s, annealing at 63°C for 30 s, extension at 72°C for 30 s, and a final extension of 72°C for 1 min in a thermal cycler (MX4005P). Quantitative PCR library amplification test and PCR library amplification were performed as previously described^49^. Sequencing reads were trimmed (TrimGalore v0.6.4) and aligned to the reference genome (GRCm38) using BWA (v0.7.17). Subsequent QC and filtering was performed using picard (v2.23.1), SAMtools (v1.10), BEDtools (v2.29.2), and BAMtools (v2.5.1), followed by peak calling using MACS2 (v2.2.7.1) and annotation via HOMER (v4.11). Statistically significant differential peaks were identified between groups using DESeq2 (v1.26) and visualized using R.

### Seahorse Assay for metabolic activity

*Cx3*cr1^CreERT2-IRES-YFP/+^Rosa26^floxed-tdTomato/+^ and CX3CR1-cre mice (control group) received tamoxifen at one day before and one day after Nb inoculation (-1, +1 day) and were sacrificed at day 7. Lung single cells were isolated and td tomato+ and td tomato-alveolar macrophages from reporters or alveolar macrophages from control group mice were sorted via flow cytometry. Alveolar macrophages (∼75,000 cells/well) from each group were seeded into CellTak (Corning, # 354240) coated XFp plates in XF RPMI complete assay media, and incubated at 37°C without CO_2_ for 45 minutes. The assay was run on a Seahorse XFp Extracellular Flux Analyzer with each group of AMs run in duplicate. After 20 cycles of basal measurement, Rotenone and Antimycin A (Seahorse XF Cell Mito Stress Kit), inhibitors of mitochondrial complex I and III, respectively, were injected into each well for a final concentration of 0.5 uM/well. Oxidative phosphorylation was measured by the oxygen consumption rate (OCAR; pmol/min) while glycolysis was measured by the extracellular acidification rate (ECAR; mpH/min) due to the accumulation of lactate or other metabolic acids and release of protons into the extracellular medium.

### 5-Ethynyl-2′-deoxyuridine (EdU) incorporation assay

*Cx3*cr1^CreERT2-IRES-YFP/+^Rosa26^floxed-tdTomato/+^ mice received tamoxifen at one day before and one day after Nb inoculation (-1, +1 day), and received Edu (5mg/ml, 100 ul, i.p.) every other day starting from one day before Nb inoculation ((-1, +1, +3, +5 day) and sacrificed at day 7. Control groups received tamoxifen and Edu but were not inoculated with Nb. Lung macrophages were stained with fluorochrome-conjugated antibodies against CD45, F4/80, CD64, CD11c, Siglec-F, then Edu incorporation was assessed using Click-iT Edu flow cytometry cell proliferation assay kit (ThermoFisher Scientific, cat#C10424).

### Statistical analysis

Data were analyzed using the statistical software program Prism 8 (GraphPad Software, Inc., La Jolla, CA) and are reported as means (± SEM). Differences between two groups were assessed by student’s T-test, differences among multiple groups were assessed by one way ANOVA and individual comparisons were analyzed using Holm-Sidak test. Differences of p < 0.05 were considered statistically significant.

---

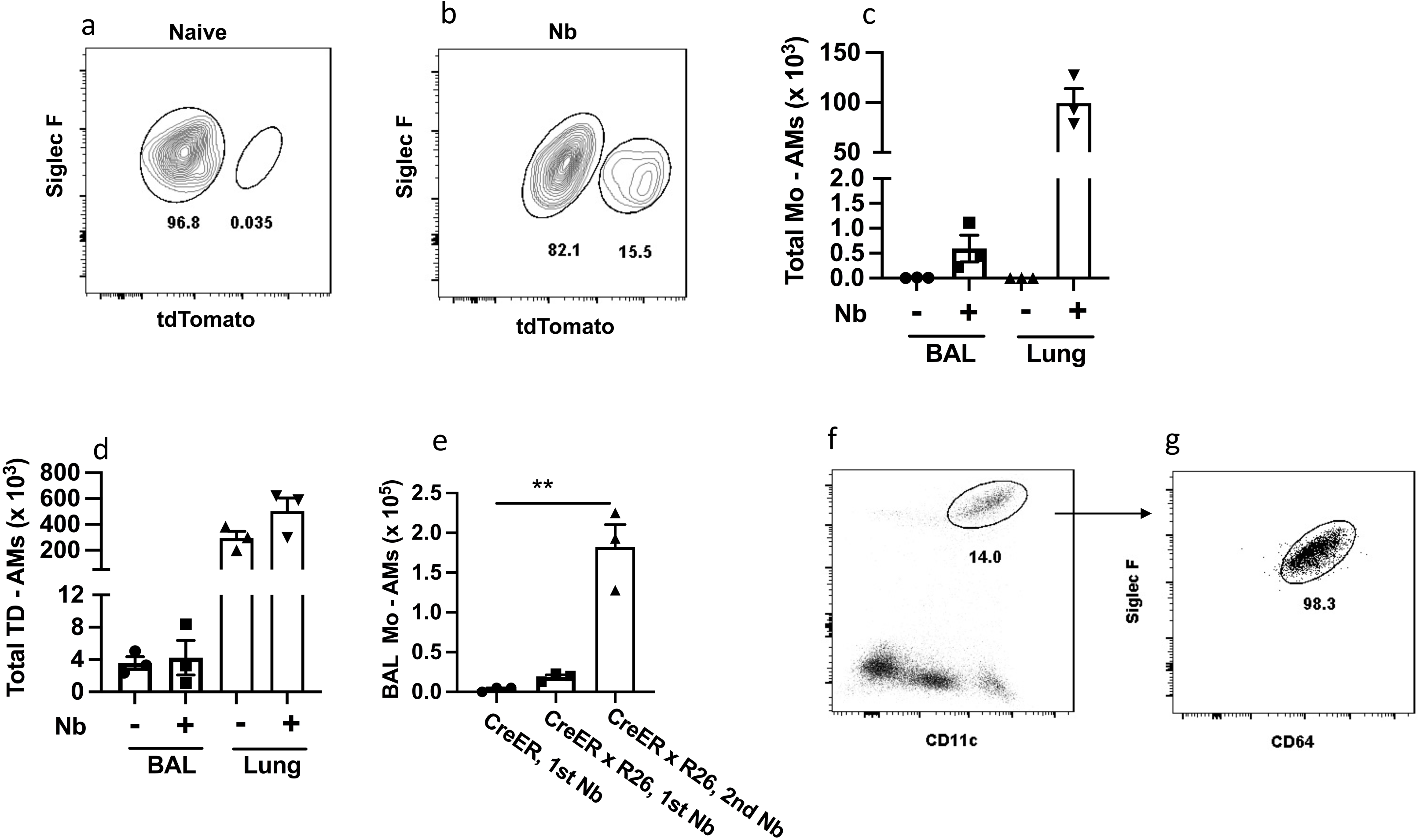

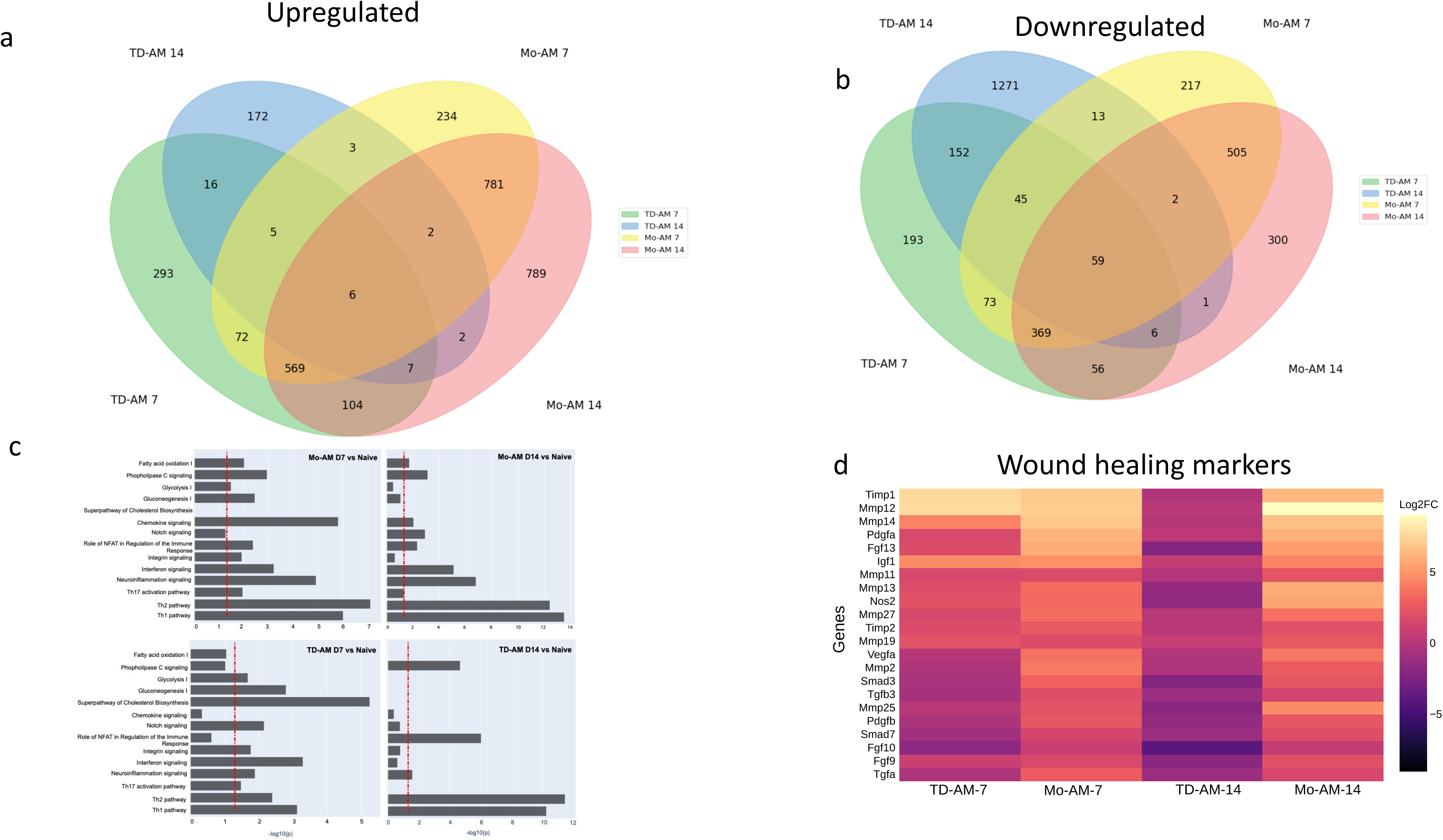

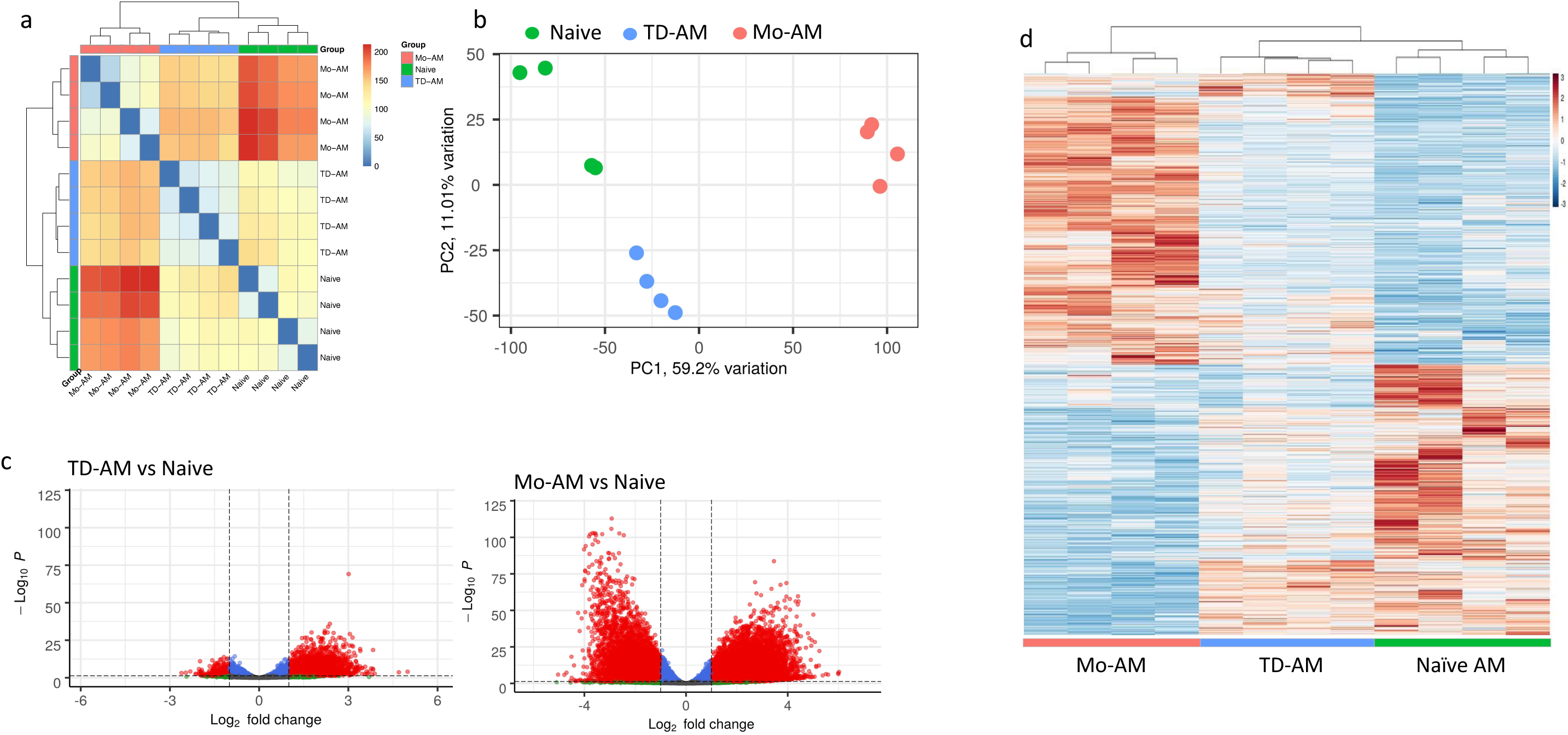

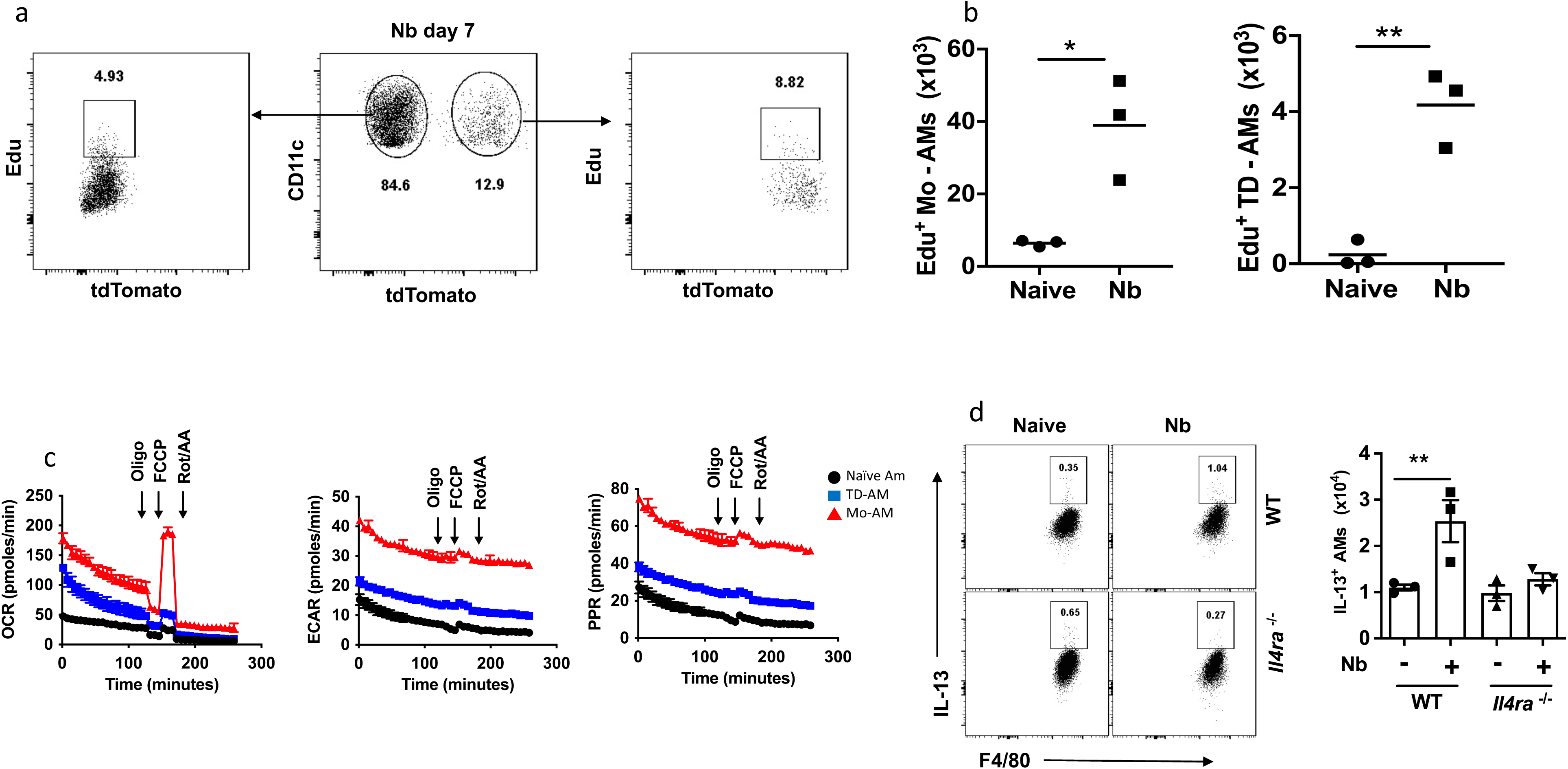

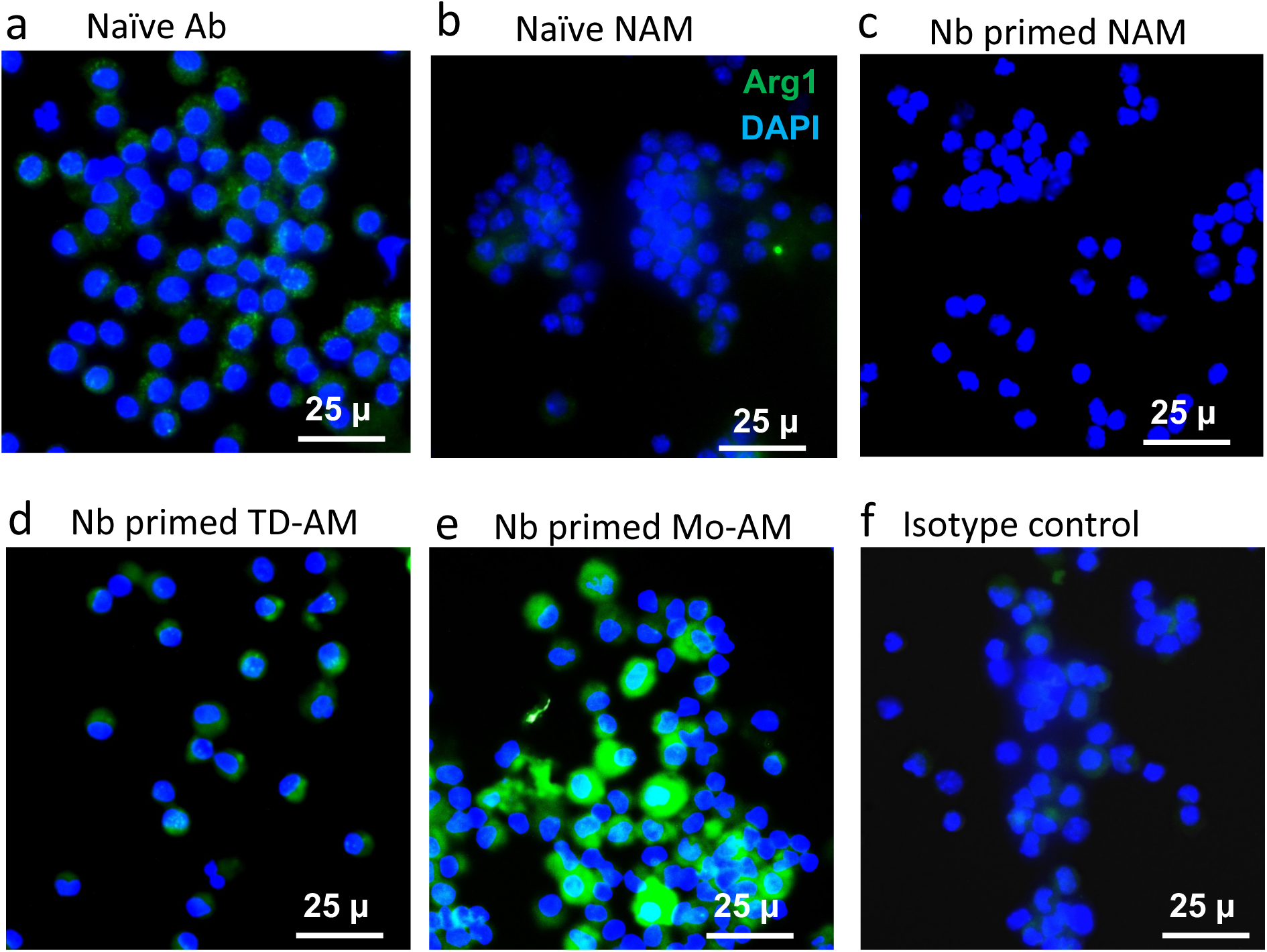

