## Supplementary Figure Legends for "Lung macrophages mediate helminth resistance through differential activation of recruited monocyte-derived alveolar macrophages and arginine depletion"

Supplementary Fig. 1 Lung tissue distribution of tissue-derived alveolar macrophages (TD-AMs) and monocyte-derived alveolar macrophages (Mo-AMs) during Nb infection.
(a,b) *Cx3*cr1$^{CreERT2-EYFP-IRES-YFP/+}$Rosa26$^{floxed-tdTomato/+}$ fate mapper mice received tamoxifen (Tam) at days -1, +1, after *N. brasiliensis* (Nb) inoculation or were uninfected. Two color contour plot is shown for Siglec-F and tdTomato expression of gated CD45$^+$,CD64$^+$, F480$^+$ cells in naïve mice (a) or in mice at day 7 after Nb inoculation (b), thereby identifying TD-AMs (Siglec-F$^+$, tdTomato$^-$) and Mo-AMs (Siglec-F$^+$, tdTomato$^+$). (c,d) Fate mapper mice were treated as above and bronchoalveolar (BAL) lavage and remaining lung tissue collected and stained for Mo-AMs or TD-AMs. (e) Fate mapper mice were administered Tam, infected with Nb and then 20 days later administered a second Nb inoculation and sacrificed 7 days later. Controls were given a primary inoculation only. (f,g) Representative flow cytometry plot of lung cell suspension at day 7 after Nb inoculation, stained as above, but showing percent of CD11c$^+$, tdTomato$^+$ cells expressing Siglec-F and CD64 (98.3%).

Supplementary Fig. 2 Gene expression profile of tdTomato$^{+/-}$ subpopulation of alveolar macrophages during N. brasiliensis infection.
*Cx3*cr1$^{CreERT2-EYFP-IRES-YFP/+}$Rosa26$^{floxed-tdTomato/+}$ reporter mice received tamoxifen at days -1, +1, after *N. brasiliensis* inoculation. At day 7 and day 14, Lung tdTomato$^+$ (Mo-AM) and tdTomato- (TD-AM) alveolar macrophages were sort-purified for RNA-seq analysis. (a-b) Venn diagram illustrating overlap of respectively upregulated or downregulated significant genes (FDR adjusted p-value <0.05) across all sample groups as compared to AMs from naïve mice. (c, d) Selected Ingenuity Pathway (IPA- Qiagen) enrichment analyses for respective sample compared to AMs from naïve mice. Dotted red bar illustrates significant enrichment cutoff (p<0.05). (d) Expression of selected characteristic wound healing response markers in TD-AMs and Mo-AMs at day 7 and day 14 after Nb inoculation as expressed relative to AMs from naïve mice ($Log_2$ fold change).

Supplementary Fig. 3 Transposase – accessible chromatin with sequencing (ATAC-seq) assay of tdTomato+/- subpopulations of alveolar macrophages after *N. brasiliensis* infection.
*Cx3*cr1$^{CreERT2-EYFP-IRES-YFP/+}$Rosa26$^{floxed-tdTomato/+}$ reporter mice received tamoxifen at days -1, and +1 after *N. brasiliensis* (Nb) inoculation. At day 7 after inoculation, tdTomato$^+$ monocyte-derived alveolar macrophages (Mo-AMs) and tdTomato$^-$ tissue-derived AMs (TD-AMs) were sorted-purified for ATAC-seq analysis and compared to naïve AMs, with 4 mice/treatment group. (a) Pairwise Euclidean distance calculation of the chromatin accessibility profiles showing TD-AM from Nb inoculated mice were more similar to naïve alveolar macrophages than Mo-AMs from Nb inoculated mice. (b) Principal component analyses of chromatin accessibility profiles of treatment groups. (c) Differential volcano plot analyses revealed greater accessibility of regulatory elements in the Mo-AMs from Nb inoculated mice. (d) Global heat map analysis of accessible regulatory elements shows distinct patterns in Mo-AMs.

Supplementary Fig. 4 Cell proliferation and metabolic analysis of lung macrophage subsets and IL-13 expression in macrophages after *N. brasiliensis* infection.
*Cx3*cr1$^{CreERT2-EYFP-IRES-YFP/+}$Rosa26$^{floxed-tdTomato/+}$ mice received tamoxifen at days -1, +1,

and Edu at days 0, +3, +5 after *N. brasiliensis* (Nb) inoculation. At day 7 after Nb inoculation, whole lung tdTomato$^{+}$ monocyte-derived alveolar macrophages (Mo-AMs) and tdTomato$^{-}$ tissue-derived alveolar macrophages (Td-AMs) were analyzed by flow cytometry for Edu incorporation. (a) a representative of flow cytometric plots showing incorporation of Edu in tdTomato$^{+}$ and tdTomato$^{-}$ AMs. (b) total Edu$^{+}$ tdTomato$^{+}$ and Edu$^{+}$tdTomato$^{-}$ AMs. Each symbol represents an individual mouse and horizontal lines indicate the mean, and data are representative of two independent experiments. *p<0.05, **p<0.01.AMs (c) *Cx3*cr1$^{CreERT2\text{-}EYFP\text{-}IRES\text{-}YFP/+}$Rosa26$^{floxed\text{-}tdTomato/+}$ were administered tamoxifen and Nb inoculated as in (a). At day 7 after inoculation, Mo-AMs, TD-AMs, and naïve alveolar macrophages were sort-purified and real-time metabolic activity assessed by Seahorse analysis. The oxygen consumption rate (OCR) and extracellular acidification rate (ECAR) were measured and inhibitors including Oligo, FCCP and Rotenone (Rot) and Antimycin A (AA) were added to block mitochondrial activity. Data shown are representative of two independent experiments. (d) Representative of flow cytometry plots (left panel) and lung macrophage numbers (right panel) for cytoplasmic IL-13 protein expression at day 7 after *N. brasiliensis*-inoculation of WT, or *Il4ra*$^{-/-}$ BALB/c mice.

Supplementary Fig. 5 Arginase 1 is preferentially expressed in lung Mo-AMs after *N. brasiliensis* inoculation.

*Cx3*cr1$^{CreERT2\text{-}EYFP\text{-}IRES\text{-}YFP/+}$Rosa26$^{floxed\text{-}tdTomato/+}$ mice received tamoxifen at days -1, +1, after *N. brasiliensis* (Nb) inoculation. At day 7 after Nb inoculation, lung cell suspensions were stained as already described for tdTomato$^{+}$ monocyte derived alveolar macrophages (Mo-AMs), tissue-derived alveolar macrophages (TD-AMs), and non-alveolar macrophages (NAMs). Cell populations were sort purified and cytospins prepared, and stained with APC-anti-Arg1 Ab. (a) naïve AM, (b) naïve NAM, (c) Nb primed NAM, (d) Nb primed TD-AM, (e) Nb primed Mo-AM, (f) Isotype control staining for Nb primed Mo-AM. Image is representative of two independent experiments.
